# CRAGE-RB-PI-seq enables transcriptional profiling of rhizobacteria during plant-root colonization

**DOI:** 10.1101/2024.11.19.624340

**Authors:** Tomoya Honda, Sora Yu, Dung Mai, Leo Baumgart, Gyorgy Babnigg, Yasuo Yoshikuni

## Abstract

Plant roots release a wide array of metabolites into the rhizosphere, shaping microbial communities and their functions. While metagenomics has expanded our understanding of these communities, little is known about the physiology of their members in host environments. Transcriptome analysis via RNA sequencing is a common approach to learning more, but its use has been challenging because plant RNA masks bacterial transcripts. To overcome this, we developed randomly-barcoded promoter-library insertion sequencing (RB-PI-seq) and combined it with chassis-independent recombinase assisted genome engineering (CRAGE), using *Pseudomonas simiae* WCS417 as a model rhizobacterium. This method enables targeted amplification of barcoded transcripts, bypassing plant RNA interference and allowing measurement of thousands of promoter activities during root colonization. Our analysis revealed time-resolved dynamics of promoter activities, highlighting early transcriptional reprogramming as a key determinant of successful colonization. Additionally, we discovered that transcriptional activation of xanthine dehydrogenase and a lysozyme inhibitor are crucial for evading plant immune system defenses. This framework is scalable to other bacterial species and provides new opportunities for understanding rhizobacterial gene regulation in native environments.

## Introduction

Plant roots secrete nutrient-rich compounds into the surrounding soil, creating unique habitats that support diverse microbial communities^1,2^. These microbial communities, in turn, play critical roles in plant physiology by promoting growth, protecting against disease, or, in some cases, causing infection^3–5^. Over the past decade, extensive metagenomic studies have identified the members of microbial communities across various plant species, genotypes, and growth environments^6–10^. The next major challenge is to develop a framework that predicts microbial gene functions, regulation, and impacts on plant physiology. Achieving this requires in-depth knowledge of the physiology of individual microbial species within their host environments. In particular, studying key organisms such as plant-growth-promoting rhizobacteria and pathogens is critical for advancing sustainable agricultural systems.

Among plant-associated bacteria, *Pseudomonas* species have been extensively studied because of their roles in promoting plant growth and protecting against disease. In particular, *Pseudomonas simiae* WCS417 has served as a model bacterium, with traits beneficial to its hosts and a remarkable ability to colonize a wide range of plant species^11^. Previous studies have uncovered molecular mechanisms underlying these symbiotic relationships, such as siderophore production^12^ and modulation of plant immune systems^13,14^. Additionally, transposon mutagenesis sequencing of *P. simiae* WCS417 identified genetic determinants that influence early-stage root colonization, including through chemotaxis and substrate utilization^15^. These findings suggest that bacteria engage in sophisticated chemical interactions with host plants during colonization. However, global profiling of these physiological adaptations remains limited.

Transcriptomics is a powerful tool for characterizing the temporal dynamics of global gene expression. In host plants, transcriptomic studies have revealed coordinated cascades of gene expression in response to microbial colonization or microbe-associated molecular patterns^16–18^. Although the host responses have been studied at both population and single-cell levels, the adaptations of rhizobacteria remain less explored. This gap is primarily due to technical challenges, as bacterial RNA is significantly outnumbered by host-plant RNA. This issue is common in plant-associated bacteria, and a similar issue is found in bacteria associated with other hosts^19–22^. To address this, various strategies have been explored, including deep sequencing^23,24^, depletion of host and bacterial rRNA^23–25^, enrichment of bacterial mRNA using customized probes^22,26^, and physical separation of bacterial cells from host tissues through techniques such as laser microdissection^27^, flow cytometry^28^, and density-gradient centrifugation^26^. Alternatively, in vitro cell culturing with root exudates has been a common approach. Although the choice of approach depends on the research question, profiling transcriptomes in planta with high efficiency and scalability remains the most desirable option.

Synthetic biology offers an alternative way to profile global transcriptional activities. Using DNA barcodes to report promoter activity enables quantification through a straightforward workflow involving PCR amplification and barcode sequencing. This method has been successfully used in model prokaryotic and eukaryotic organisms, revealing unique features of promoter regulatory elements^29–32^. In the present study, we introduced a DNA-barcoded promoter library into *P. simiae* WCS417, using chassis-independent recombinase-assisted genome engineering (CRAGE)^33,34^, a versatile strain-engineering technology developed in our group. We then established an experimental workflow to characterize in planta promoter activities by selective amplification of barcoded transcripts, overcoming interference from plant RNA. We term this workflow as “CRAGE randomly-barcoded promoter-library insertion sequencing (CRAGE-RB-PI-seq),” or PI-seq in short. Using this method, we identified a set of uniquely regulated promoters involved in plant root colonization and validated their activities and functions through a series of independent experiments. The framework we developed enables rapid characterization of transcriptional activities and can be applied to a wide range of bacterial species, providing new insights into gene function and regulation in native environments.

## Results

### Development of the CRAGE-RB-PI-seq workflow

To generate a barcoded promoter library, we first computationally extracted the 140 bp upstream sequences from the start codons of all annotated protein-coding sequences in the *P. simiae* WCS417 genome (**Fig.1 A**). We chose a length of 140 bp, as this length accommodates the 66 % of transcriptional factor binding sites in *P. simiae* WCS417 (**Fig.S1**), and the typical 20-80 bp distance of transcription start site in bacteria^29,35–37^, while also being feasible to acquire oligo pool from Twist Biosciences at the time. To optimize versatility, we divided promoter libraries into three groups according to the distances to upstream genes: group I (≤30 bp), group II (31–139 bp), and group III (≥140 bp). This grouping later helped us reduce library complexity by selecting groups with active promoters, as the presence of transcriptional start sites (TSSs) generally correlate with intergenic length^38^. Each library was synthesized, barcoded, and cloned into a CRAGE accessory vector (**Fig.1B**), then transformed into a conjugal *E. coli* donor strain.

**Figure 1:**
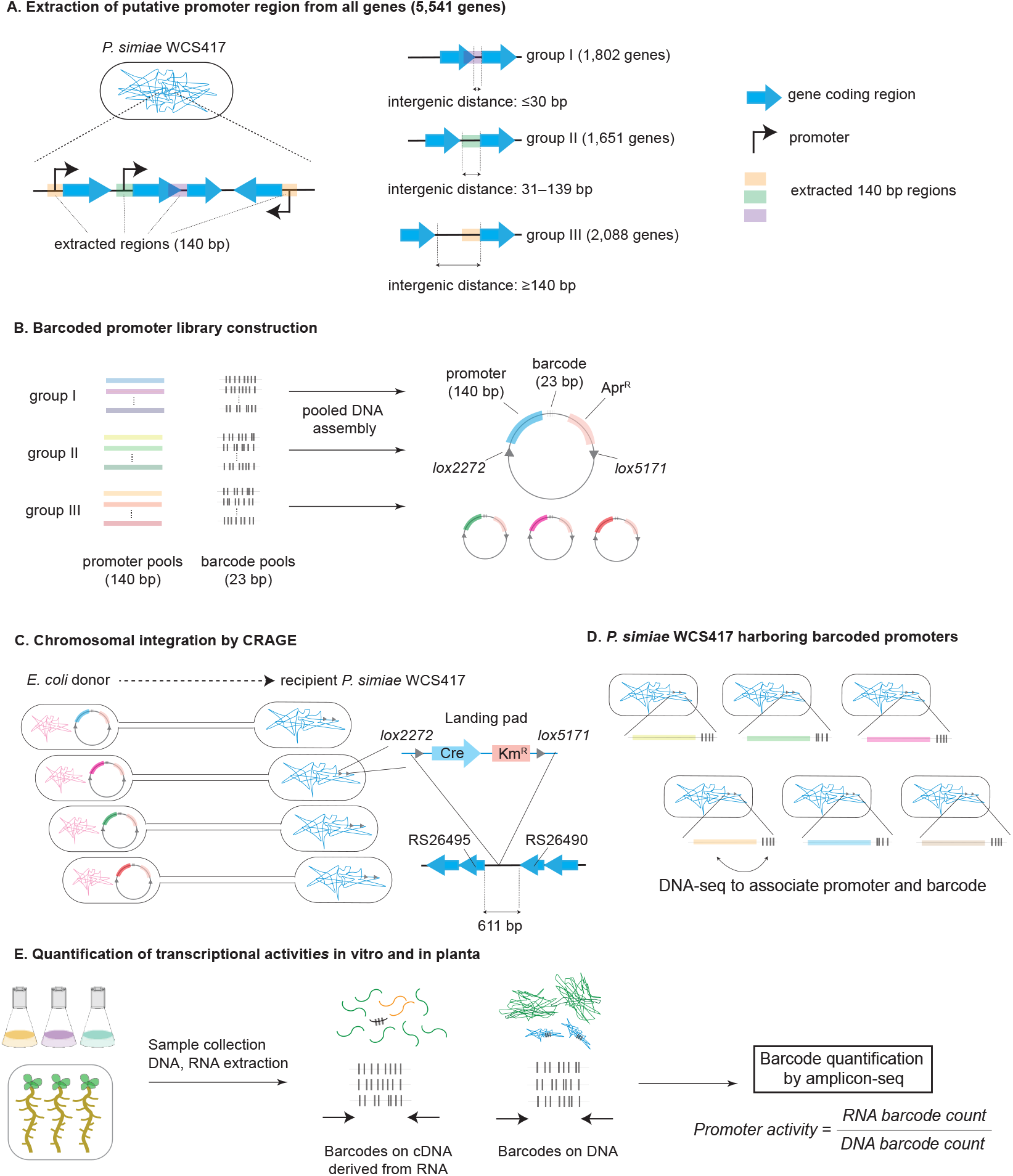
Development of the CRAGE-PI-seq workflow. (**A**) A total of 5,541 sequences, each 140 bp sequences immediately upstream of start codons, were computationally extracted. These promoter sequences were categorized into three groups according to the distances to upstream genes: group I (≤30 bp), group II (31–139 bp), and group III (≥140 bp). (**B**) Each promoter library was synthesized, barcoded with 23 bp sequences, and cloned into the CRAGE accessory vector (pW26_mod), which included an apramycin-resistant marker (Apr^R^) flanked by two mutually exclusive lox sites (*lox2272* and *lox5171*), necessary for recombination. (**C**) The plasmids were transformed into an *E. coli* donor strain and conjugated into *P. simiae* WCS417 (strain SB599), which harbored a CRAGE landing pad on the chromosome. The landing pad is located between locus tags PS417_RS26490 and PS417_RS26495, without disrupting any coding sequences^34^. Cre-mediated recombination integrated the barcoded promoters into the landing pad. (**D**) The promoter-barcode regions were amplified from the genomic DNA and sequenced via targeted DNA-seq to associate each promoter with its corresponding unique barcodes. (**E**) The transcriptional activity of promoters was characterized by growing cells under various conditions, extracting DNA and RNA, and amplifying barcode regions from the DNA and cDNA. The cDNA was synthesized from RNA by selective reverse transcription. Promoter activities were then quantified by normalizing RNA barcode counts to DNA barcode counts for each gene.

The promoter libraries were chromosomally integrated into *P. simiae* WCS417 (strain SB599) using CRAGE (**Fig.1C**). The integration site lies in the middle of an intergenic region, without disrupting any genes^34^ (at nucleotide location 268,402 between locus tag PS417_RS26490 and 26495), and our extended analysis confirmed that the engineering process did not alter cellular physiology (**Fig.S2-3**). DNA sequencing (DNA-seq) was used to link each promoter with unique barcodes (**Fig.1D**). This process successfully cloned 91% of promoters (5,040 out of 5,541), each tagged with unique barcodes (**Supplementary Data 1)**. Pooled cell libraries were cultured under various conditions, both in vitro and in planta (**Fig.1E**). Promoter activities were quantified using targeted RNA-seq and DNA-seq, normalizing RNA-derived barcode counts to DNA-derived counts (**Fig.1E**).

### Transcriptional profiling of *P. simiae* WCS417 under various in vitro conditions

To test the functionality of our promoter library, we first grew the pooled library (groups I–III) under five different conditions, preparing sequencing samples for both PI-seq and conventional RNA seq (**Fig.2A**). PI-seq consistently demonstrated high library coverage and reproducibility across conditions. For example, DNA barcode analysis showed that under the reference condition (20 mM glucose), 98.8% of promoters (4,982 out of 5,040 in the library) were covered at least 50 times per million reads, and the histogram showed a narrow distribution (**Fig.2B**). The data reproducibility between biological replicates was high (Pearson’s r > 0.99) (**Fig.2C**), indicating sufficient library coverage under these culture conditions.

**Figure 2:**
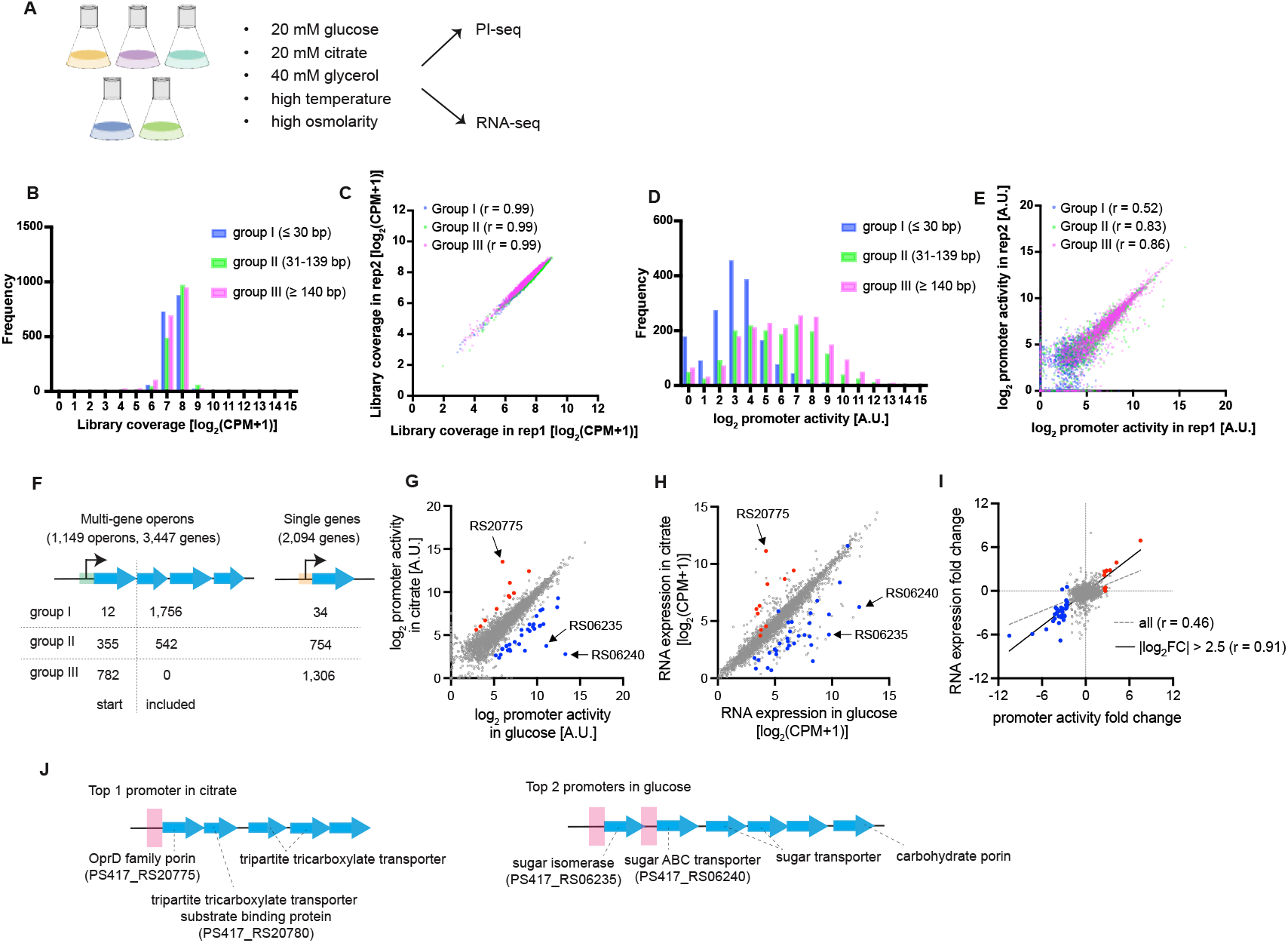
Characterization of promoter activities in liquid cultures. (**A**) Experimental overview. A pooled library of groups I–III was grown under five different conditions. PI-seq and RNA-seq libraries were created for each condition (*n* = 2 biological replicates). (**B-E**) Analysis of PI-seq data for the cells grown in 20 mM glucose, used as an example. DNA barcode counts showed a narrow distribution and high coverage of the designed library (B). Comparison of the library coverage for two biological replicates showed high reproducibility (C). In contrast, RNA barcode counts normalized to DNA barcode counts showed a large variation in promoter activity. Groups II and III promoters (green and magenta bars, intergenic distance ≥ 31 bp) were more active than group I promoters (blue bars, intergenic distance ≤ 30 bp) (D). The biological replicates for groups II and III showed high reproducibility of promoter activity, while the reproducibility of group I promoter activity was weaker (E). Pearson’s r correlation values for each group are shown in parentheses. (**F**) In silico prediction of operon structure using Rockhopper identified 1,149 operons and 2,094 single genes. From these 3,243 genes, 2,880 promoters were successfully cloned into the library and used to compare PI-seq and RNA-seq results. (**G**) Comparison of promoter activity under two conditions. PI-seq data showing promoter activity under 20 mM glucose (x-axis; blue points) and under 20 mM citrate (y-axis; red points) highlights upregulated promoters. Differentially activated promoters were selected based on fold changes (FCs) (log_2_FC > 2.5) after filtering out low-activity promoters (log_2_ promoter activity < 2). (**H**) Comparison of gene expression under 20 mM glucose and 20 mM citrate conditions using RNA-seq data. Genes driven by the promoters identified by PI-seq in (G) are highlighted in the same colors. (**I**) Correlation of expression changes between PI-seq and RNA-seq. Comparison of fold changes in expression between PI-seq (x-axis) and RNA-seq (y-axis) shows strong quantitative agreement for the promoters with large fold changes (log_2_FC > 2.5), indicated by red and blue points. The fold changes were calculated by analyzing the expression under the 20 mM citrate condition divided by that under the 20 mM glucose condition for each method. Pearson’s r correlation values are provided (all genes: dashed gray line; genes with log_2_FC > 2.5: solid black line). (**J**) Illustrations of the most upregulated promoters and their associated downstream genes. Consistent with the experimental conditions, the promoter most upregulated under 20 mM citrate conditions drives genes encoding porin and a tripartite tricarboxylate transporter, while the two promoters most upregulated under 20 mM glucose conditions drive sugar transporter genes within the same operon. The 140 bp promoter regions are highlighted in pink.

Promoter activity — determined by normalizing RNA barcode counts to DNA barcode counts — showed a large variation within and between groups I–III, with the activity levels differing by several orders of magnitude (**Fig.2D**). In particular, group II and III promoters (green and magenta bars in the figure) exhibited overall higher activity than group I promoters (blue bars). The results of biological replicates were highly reproducible for group II and III libraries (Pearson’s r > 0.83), while the results between biological replicates for group I were less reproducible (Pearson’s r ∼ 0.52) (**Fig.2E**), a trend noted across all growth conditions (**Fig.S4**).

We hypothesized that the lower activity of group I promoters (derived from ≤30 bp intergenic regions) might be because many promoters were part of operons, and therefore lacked active transcription start sites^38^. To investigate this, we used Rockhopper^39^ to predict operon structures in silico, incorporating *P. simiae* WCS417 genome sequence data and RNA-seq data. This analysis identified 1,149 operons comprising 3,447 genes, as well as 2,094 single isolated genes (**Fig.2F, Supplementary Data 2**). As expected, most group I promoters were part of operons, while all group III promoters corresponded to either operon start sites or isolated genes. Of the 3,243 genes (1,149 operonic + 2,094 single), 2,880 promoters were successfully cloned into our libraries. Subsequent analysis focused on these 2,880 promoters to further validate their activities.

### Analysis of differentially induced promoters from PI-seq data

We next compared PI-seq data from cells grown under 20 mM glucose conditions with the data from 20 mM citrate conditions. These two carbon sources require distinct transporters and metabolic pathways^40^, allowing us to assess the promoter library’s performance. Differential analysis identified 32 promoters that were uniquely upregulated under glucose conditions, and 10 promoters that were uniquely upregulated under citrate conditions (**Fig.2G**). RNA expression driven by these upregulated promoters were similarly upregulated, as RNA-seq verified these patterns (**Fig.2H**). Comparison of fold changes in promoter activity and RNA expression demonstrated a strong quantitative agreement between PI-seq and RNA-seq, as illustrated in **Fig.2I**. Notably, there was a strong correlation between the two methods for differentially induced promoters (**Fig.2I**, blue and red points, fitted with a solid line: Pearson’s r = 0.91), although some fraction of promoters did not match RNA-seq results, which may reflect missing regulatory elements (**Fig.S1**) or post-transcriptional regulation^41,42^ in our library design.

Additional analysis of other culture conditions confirmed similar trends for both PI-seq and RNA seq (**Fig.S5**). Unsupervised clustering analysis of the PI-seq data revealed distinct clustering depending on culture conditions, consistent with the pattern in RNA-seq data (**Fig.S6**). Importantly, PI-seq captured physiologically relevant changes in promoter activity (**Fig.2J**). For example, under glucose conditions, the two most upregulated promoters (annotated by arrows in **Fig.2G**) drive genes (PS417_RS06235, PS417_RS06240) involved in glucose binding and transport, while under citrate conditions, the most upregulated promoter drives an operon that encodes porin and tripartite-tricarboxylate transporter-substrate-binding protein (PS417_RS20775, PS417_RS20780), both functional for citrate uptake^43,44^. These findings align with expected differences corresponding to the different culture conditions, and the large fold changes in RNA expression were confirmed by RNA-seq (annotated in **Fig.2H**). Overall, these analyses suggest that PI-seq is effective at identifying a core set of promoters that are reprogrammed in response to environmental changes.

### Transcriptional profiling of *P. simiae* WCS417 during Arabidopsis root colonization

Using the engineered strains, we investigated *P. simiae* WCS417 promoter activity during colonization of Arabidopsis roots (**Fig.3A**). Pooled promoter libraries from groups II and III were grown in M9–glucose medium and inoculated with 10-day-old seedlings on phytagel plates supplemented with 0.5X Murashige & Skoog (MS) basal salts. Group I promoters were omitted because of their low activity. 10-day-old seedlings were chosen, because this developmental stage is commonly used in other agar-based root colonization studies^4,15,45^. Whole root samples (∼15 seedlings per sample) were collected at 10 minutes, 3 hours, 24 hours, and up to 7 days. Promoter activity was quantified by barcode counts from extracted DNA and RNA. Despite having ∼100 times fewer cells in planta (∼2×10^7^) than the number of cells in liquid cultures (1.6×10^9^) and handling samples contaminated with ∼20 mg of plant roots, we achieved high library coverage and reproducible results (**Fig.3B–E**). This demonstrates the utility of our approach, as conventional RNA-seq often requires deep sequencing to overcome plant RNA contamination and obtain sufficient bacterial reads (**Table S2**). We furthermore confirmed that the PI-seq library strain elicited similar plant transcriptional responses as the wild-type *P. simiae* WCS417 (**Fig.S7**).

**Figure 3:**
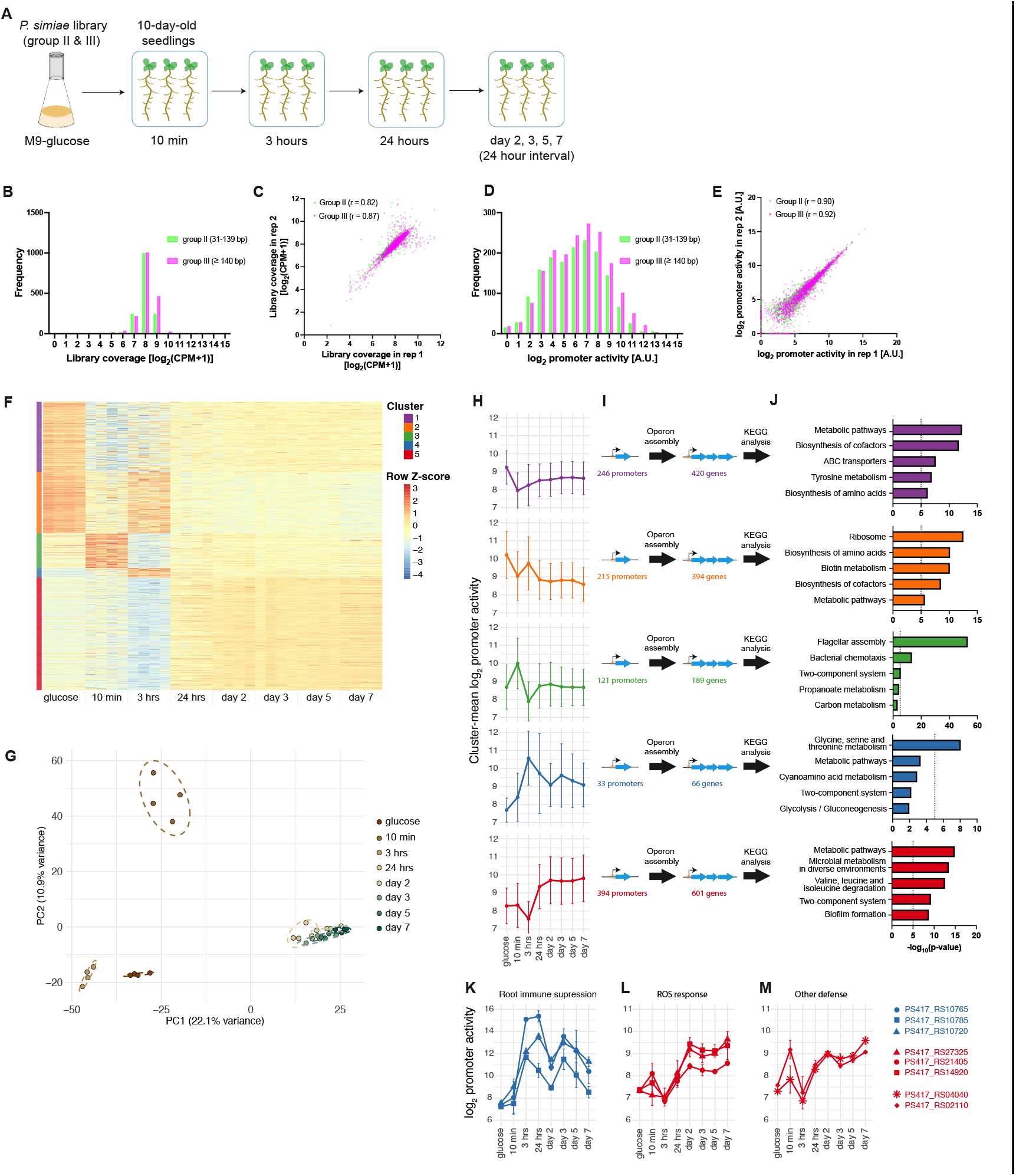
Characterization of promoter activities during Arabidopsis root colonization. (**A**) *P. simiae* WCS417 populations containing promoter libraries from groups II and III were grown in M9-glucose medium and inoculated onto phytagel plates with 10-day-old *Arabidopsis thaliana* seedlings. Group I was excluded because of low promoter activity (Fig.2). Whole root samples were collected at 10 minutes, 3 hours, 24 hours and up to 7 days. Barcode sequencing libraries were prepared by targeted PCR amplification from extracted RNA and DNA to assess promoter activity (*n* = 4 biological replicates per time point). (**B–E**) PI-seq data from root colonizing cells (day-7) demonstrated a high library coverage (B) and reproducibility (C), a large variation (D) and high reproducibility of promoter activity across biological replicates (E). (**F**) Promoters differentially expressed relative to the glucose condition were identified (|log_2_FC| > 1 and BH-adjusted p-value < 0.05) with a total of 1,009 promoters. Log_2_ promoter activity was z-score normalized per promoter and visualized as a heat map. Rows were hierarchically clustered and split into five clusters based on their activity profiles. (**G**) Principal component analysis of promoter activity across biological replicates. Points are colored by the sampling point. (**H**) Cluster-mean log_2_ promoter activity values ± SD were plotted across sampling points. Colors match cluster assignments in the heat map. (**I**) Predicted operon structures (from Fig.2F) were linked to promoters, and the corresponding genes were subjected to KEGG pathway analysis. (**J**) Top-5 enriched KEGG pathways in each cluster. Dashed lines indicate statistical significance thresholds (adjusted p-value < 10^−5^). (**K-M**) Examples of promoter activities: (K) plant immune suppression genes in cluster 4, and (L–M) ROS response and other defense related genes in cluster 5.

To understand colonization dynamics, we compared promoter activities at each sampling point against the M9–glucose condition and identified a total of 1,009 differentially expressed promoters. These promoters were grouped into five clusters by hierarchical clustering based on their activity profiles (**Fig.3F**). Promoter activities changed most dynamically within the first 24 hours, after which they remained relatively stable. This pattern was also evident in principal component analysis (**Fig.3G**). Each cluster displayed distinct temporal patterns of activation and inactivation (**Fig.3H**). Promoters in clusters 1 and 2 were highly active in M9–glucose, although cluster 2 exhibited a transient reactivation at 3 hours. Cluster 3 and cluster 4 showed transient activation at 10 minutes and 3 hours, respectively. Cluster 5 displayed a gradual increase in activity after 24 hours, marking the late stage of colonization.

We next examined the genes driven by these promoters using Rockhopper operon structure analysis (**Fig.3I, Supplementary Data 4**). Functional insights from KEGG pathway enrichment revealed distinct pathways in each cluster (**Fig.3J**). Cluster 3 was enriched for flagellar assembly and bacterial chemotaxis, whose strong induction as early as 10 minutes implies their importance in early root colonization, consistent with previous reports^15,46,47^. Cluster 4 was enriched for glycine, serine, and threonine metabolism. Its activation coincided with cluster 2, which included ribosome and amino acid biosynthesis pathways, suggesting that the 3-hour induction reflects initiation of cell growth by taking up root exudates. Cluster 1 was enriched for ABC transporters and biosynthesis of cofactors and amino acids, consistent with their strong expression only in M9–glucose. In contrast, cluster 5 was enriched for microbial metabolism in diverse environments, two-component systems, and biofilm formation. The functional distinction between cluster 5 and clusters 1-2, which were dominated by biomass production pathways such as ribosome and amino acid biosynthesis, suggests a physiological shift from a metabolically active state to more adhesive and protective states during colonization, in line with prior studies^23,48,49^. Finally, the detailed analysis of the top-enriched “metabolic pathways” in clusters 1 and 5 revealed activation of distinct metabolic programs engaged under each condition (**Fig.S8**).

Additionally, several promoters linked to genes essential for root colonization were upregulated. Cluster 4 included pyrroloquinoline quinone (PQQ) dependent genes (PS417_RS10765, PS417_RS10785, PS417_RS10720) that suppress root immune responses via gluconic acid production^14^ (**Fig.3K**). Cluster 5 contained catalase (PS417_RS27325) and DNA repair enzymes (PS417_RS21405, PS417_RS14920), both involved in mitigating reactive oxygen species (ROS) likely produced by plants as a defense response^50,51^ (**Fig.3L**). Other upregulated promoters in cluster 5, potentially important for colonization, included those driving a xanthine dehydrogenase (PS417_RS0404) and a hypothetical protein (PS417_RS02110) predicted to be a member of the MliC (membrane-bound lysozyme inhibitor of c-type lysozyme) family protein (**Fig.3M**); we experimentally validated the latter two functions, as shown in **Fig.4**.

**Figure 4:**
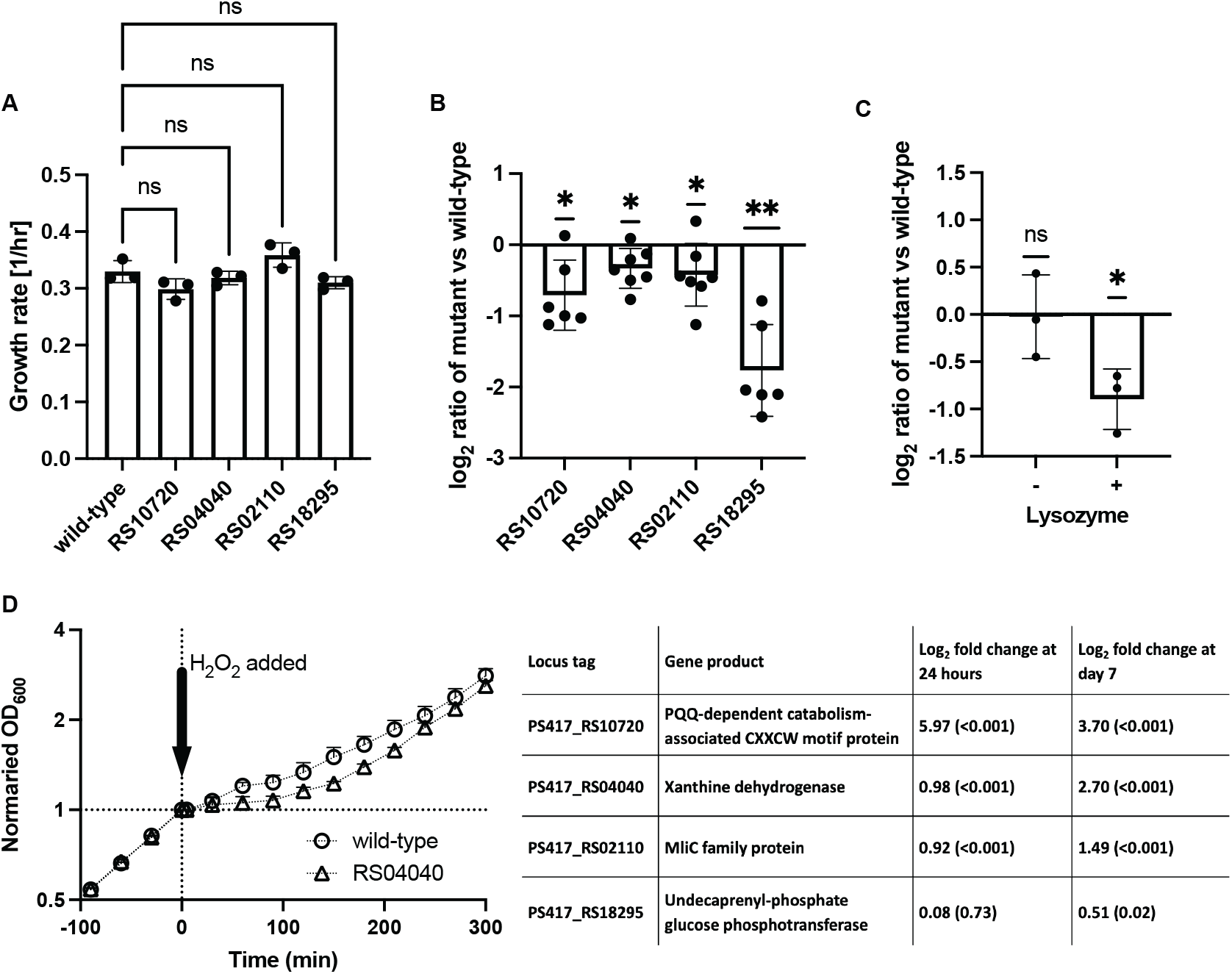
Mutant phenotypes in root colonization and stress tolerance. (**A**) Growth rates of the wild-type strain and four insertion mutants were compared in M9 media supplemented with 20 mM glucose. The growth rates of mutant strains were indistinguishable from those of the wild-type strain. Statistical comparisons were performed using one-way ANOVA, comparing each mutant to wild-type. Mutants are identified by their locus tags (with “PS417_” omitted; see the legend table for full tag). (**B**) Root colonization competition assay with Arabidopsis seedlings. Wild-type and individual mutant strains were mixed 1:1 and applied to plates containing 10-day-old Arabidopsis seedlings. After 5 days, root samples were collected, and wild-type and mutant strain colonies were counted (see **Supplementary Information 5.1**). The log_2_ ratios of mutant to wild-type colonies are shown in the graph. All tested mutants were outcompeted by the wild-type strain, as indicated by their ratios falling below the line at y = 0. The bars represent mean ± SD. Statistical significance was assessed using a one-sample t-test against 0. (*, p < 0.05; **, p < 0.01). (**C**) Lysozyme assay of the wild type strain and an insertion mutant in the MliC family protein. Cultures were adjusted to OD600= 0.05 and incubated in PBS for 24 hours, either without (left) or with (right) 25 mg/mL lysozyme. Cell numbers were determined by CFU counts, and the mutant strain displayed fewer cells than the wild-type in the presence of lysozyme. (**D**) Oxidative stress tolerance assay to examine the function of xanthine dehydrogenase (PS417_RS04040). Wild-type and mutant strains were grown in M9 media supplemented with 20 mM glucose and challenged with 5 mM H_2_O_2_ at time 0. The mutant required approximately 60 minutes longer than the wild-type to restore exponential growth after oxidative stress exposure (*n* = 3 biological replicates). The legend table lists the mutant locus tags and their corresponding gene products. Log2fold changes in promoter activity at 24 hours and day 7, relative to the M9-glucose condition, are shown based on PI-seq data. Adjusted p-values are provided in parentheses.

### Characterization of mutant phenotypes

To examine the functions of genes driven by upregulated promoters, we obtained transposon insertion mutants for three genes generated by Cole *et al*^15^ (see **Fig.4** legend table) and characterized their metabolic and root colonization capabilities. In addition, we included a gene involved in polysaccharide-mediated biofilm production (PS417_RS18295), whose promoter was not identified in the differential expression analysis, although the pathway is enriched in cluster 5, for further phenotypic testing. When grown individually, all mutants displayed growth rates comparable to the wild-type strain in M9-glucose minimal medium (**Fig.4A**). However, in competitive root colonization assays, the wild-type strain significantly outcompeted the mutants (**Fig.4B**). The reduced colonization observed for mutants of the PQQ-dependent catabolism associated protein (PS417_RS10720) and the undecaprenyl-phosphate glucose phosphotransferase (PS417_RS18295) is consistent with previous findings that highlight the importance of plant immune suppression^14^ and biofilm formation^52,53^, respectively, in root colonization.

The wild-type strain also outcompeted two other mutants, carrying insertion mutations in genes PS417_RS02110 and PS417_RS04040 whose roles in the root colonization ability of *P. simiae* WCS417 had not been previously reported. According to RefSeq annotation PS417_RS02110 encodes an MliC family protein. The MliC protein from *Pseudomonas aeruginosa* and *Escherichia coli* is known to function as a c-type lysozyme inhibitor, aiding in host colonization of humans and other animals^54,55^. Although *P. simiae* WCS417’s MliC shares limited protein sequence identity with these homologs, AlphaFold 3 predicted significant structural similarities^56^ (**Fig.S9**), suggesting a similar lysozyme-inhibiting function. This is especially relevant given recent studies showing that lysozyme-like hydrolases are produced by Arabidopsis^57^ and epiphytic fungi^58^, potentially influencing microbial communities. While *Pseudomonas* species are generally lysozyme-resistant, the role of MliC may be critical in the presence of agents that destabilize the outer membrane of the bacterium, thereby increasing its susceptibility to external lysozymes^59,60^. To investigate whether MliC increases lysozyme tolerance in *P simiae* WCS417, we examined the effect of lysozyme on the wild-type strain and a mutant carrying an insertion in *mliC*. The mutant showed reduced cell numbers compared to the wild-type in the presence of lysozyme, whereas no difference was observed between them in the absence of lysozyme (**Fig.4C**), indicating that MliC contributes to *P. simiae* WCS417’s protection against lysozymes from plants and from other microbes.

The second gene, PS417_RS04040, encodes xanthine dehydrogenase (XDH), whose transcriptional activation has been linked to ROS tolerance and host infection in *Ralstonia solanacearum*^61^ and in *Borrelia burgdorferi*^62^. Since ROS generation is a common plant immune response, we hypothesized that XDH in *P. simiae* WCS417 plays a role in neutralizing plant generated ROS. To test this, we examined the effect of ROS stress on the wild-type strain and a mutant carrying an insertion in *xdh* during exponential growth in M9 minimal medium with glucose. Upon exposure to 5 mM hydrogen peroxide, both wild-type and mutant strains experienced growth arrest, while the mutant showed a longer arrest by ∼60 minutes compared to the wild-type strain (**Fig.4D**), supporting our hypothesis. These findings suggest XDH may contribute to root colonization by working in concert with other ROS-responsive enzymes such as catalases and DNA ligases.

### Application of PI-seq to Arabidopsis grown in a soil-like system

To test whether PI-seq is applicable to plants grown in a more natural system, we inoculated pooled promoter libraries from groups II and III (grown in M9–glucose medium) into the rhizosphere of 10-day-old Arabidopsis seedlings cultivated in clay (particle size ∼4 mm) supplemented with 0.5× MS basal salts. Root samples were collected after 7 and 21 days, extending the colonization period compared to the phytagel plate experiments (**Fig.5A**). Clay was selected as the substrate because it resembles physical properties of soil while also permitting gentle separation from Arabidopsis roots that are fine and fragile.

**Figure 5:**
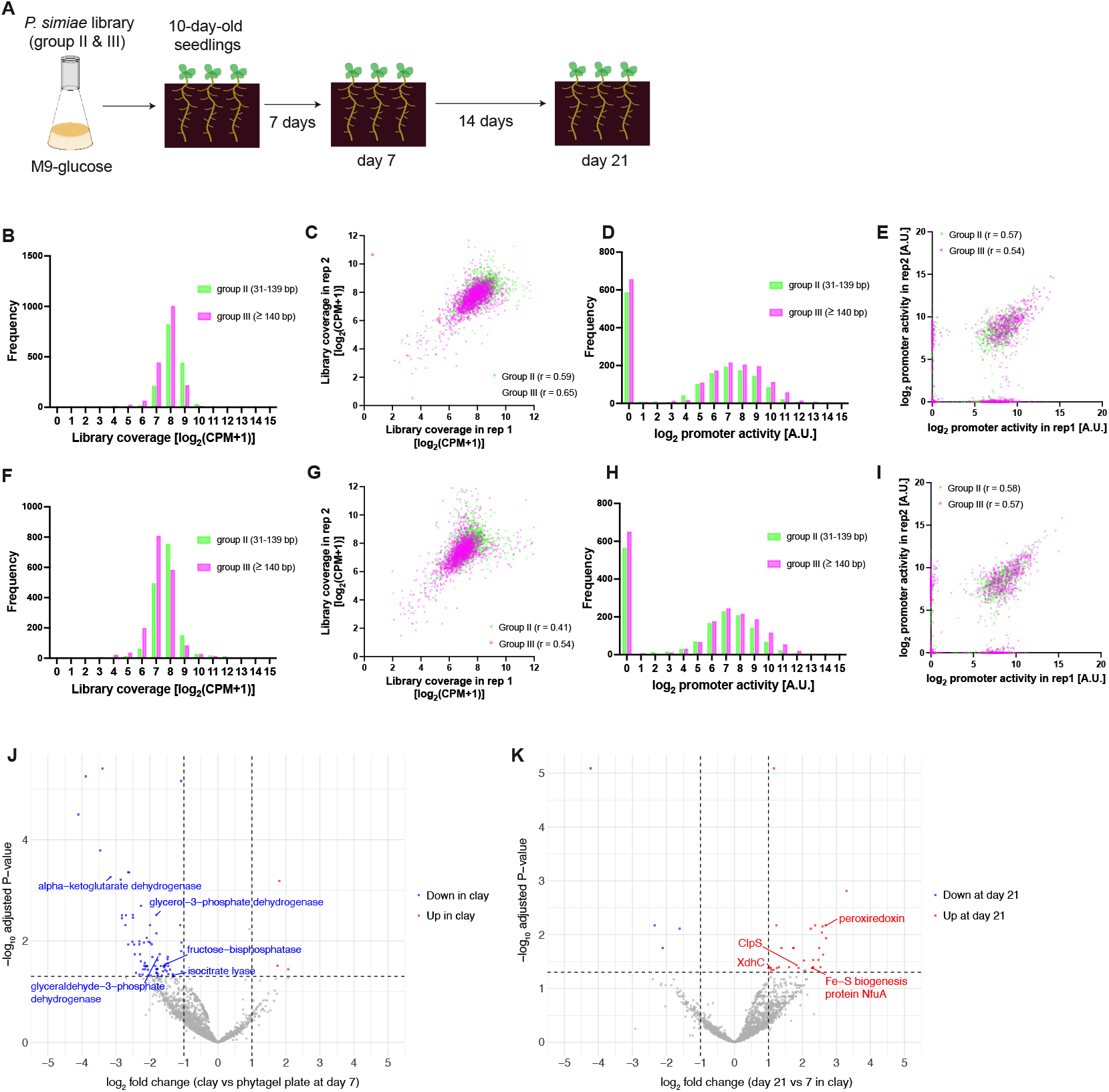
Application of PI-seq to a soil-like system. (**A**) *P. simiae* WCS417 populations containing promoter libraries from groups II and III were grown in M9–glucose medium and inoculated into clay pots with 10-day-old Arabidopsis seedlings. Root samples were collected at 7 and 21 days. Barcode sequencing libraries were prepared by targeted PCR amplification from extracted RNA and DNA to quantify promoter activity (n = 3 biological replicates per time point). (**B**) Library coverage based on DNA barcode data at day 7. (**C**) Reproducibility of DNA barcode counts across two biological replicates at day 7. (**D**) Distribution of promoter activity at day 7, showing ∼30 % inactive promoters. (**E**) Reproducibility of promoter activity across two biological replicates at day 7. (**F–I**) Same analyses as in panels (B–E), but using day 21 samples. (**J**) Volcano plot comparing promoter activity between clay and phytagel at day 7; promoters driving genes in central carbon metabolism are highlighted. (**K**) Volcano plot comparing promoter activity between day 7 and day 21 in clay; promoters driving stress response and maintenance genes are highlighted.

As in the phytagel experiments, DNA barcode analysis showed that 98 % of the 3,382 promoters in the groups II and III library were represented at least 50 times per million reads in both day 7 and 21 samples, demonstrating sufficient library coverage (**Fig.5B&F**). Biological replicates of DNA barcode counts were reproducible, although correlations were lower than those observed in phytagel plates (**Fig.5C&G**). In contrast, RNA barcodes were recovered from a subset of the library: promoter activity was detected in 70 % of promoters at day 7 and 69 % at day 21, with the remaining ∼30 % inactive (**Fig.5D&H**). This reduced activity resulted in weaker correlations across biological replicates compared to phytagel experiments (**Fig.5E&I**). This lower RNA recovery likely results from 1) adsorption of root exudates to clay particles^63^, 2) reduced nutrient availability in an open environment, and 3) diminished metabolic activity during late-stage colonization (**Fig.3**), all contributing to reduced transcriptional activity.

We next compared promoter activity at day 7 between phytagel plates and clay, filtering out promoters with zero activity in the clay dataset. Differential expression analysis identified 3 promoters upregulated and 81 downregulated in clay relative to phytagel (**Fig.5J, Supplementary Data 5**). The downregulated set in clay included genes associated with central carbon metabolism, consistent with lower metabolic activity in soil-like conditions. Comparing day 7 and day 21 in clay revealed 31 upregulated and 4 downregulated promoters at the later time point. The upregulated group included promoters driving genes for stress response and cellular maintenance such as peroxiredoxin, XDH, ClpS, and Fe-S biogenesis protein NfuA (**Fig.5K, Supplementary Data 5**), consistent with the phytagel experiments, where late-stage colonizers further shifted toward stress tolerance and long-term survival.

## Discussion

Plant-growth-promoting rhizobacteria offer a promising avenue for enhancing plant biomass and crop yields without high chemical fertilizer use. To harness their full potential, it is critical to understand their in planta physiology, including gene functions and regulatory mechanisms. However, studying these rhizobacteria in host-associated environments is challenging because of their low biomass. Traditional RNA-seq approaches for plant-colonizing bacteria often require large sample sizes and extensive rRNA depletion, followed by deep-sequencing. Even then, only about 0.5% of reads map to bacterial gene coding sequences, with a large proportion mapping to plants^23,26^ Further optimizations such as improved combinatorial rRNA removals^24^ and physical separation of cells prior to lysis^26^ are necessary to enrich bacterial mRNA when RNA-seq approaches are used.

In this study, we developed CRAGE-RB-PI-seq in the model rhizobacterium *P. simiae* WCS417 to study its transcriptional activities during Arabidopsis root colonization. This approach uses PCR amplification of RNA barcodes from a promoter library, enabling nearly 100% enrichment of bacterial transcripts. Compared with conventional deep RNA-seq, PI-seq substantially reduces sequencing depth requirements and associated reagent costs. PI-seq will also become highly effective particularly when the species of interest is present in low abundance. Although our experiment involved only ∼10^7^ cells collected from ∼20 mg of root per sample, the input cell amount could be reduced even further by increasing the number of PCR cycles. Moreover, this approach is adaptable to other bacteria compatible with CRAGE. Our current strain portfolio includes approximately 50 species across diverse phyla, many of which were isolated from soil environments. Barcoding also offers a distinct advantage in that it allows differentiation of homologous genes across different organisms; the difficulty in differentiating these is a frequent challenge in metatranscriptome analysis.

We selected the 140 bp sequences that were immediately upstream of individual genes as promoters because of the technical limitations of the oligo pools available at the time from Twist Biosciences. Despite this constraint, our results showed high correlation between PI-seq and RNA-seq, particularly in identifying differentially expressed genes (**Fig.2IJ** and **Fig.S5**). We expect that extending the promoter regions would result in even greater accuracy as it covers more transcriptional factor binding sites (**Fig.S1**); this is now possible with new oligo pools that can harbor 240 bp sequences (Twist Biosciences). Advances in oligo synthesis technologies such as enzymatic oligo synthesis and DropSynth will also allow construction of longer promoter sequences^64,65^, improving library quality. Additionally, results could be enhanced by refining library design to capture entire promoter regions, including regulatory elements and transcriptional start sites, which could be facilitated by 5’-enriched RNA-seq methods such as differential RNA-seq^66^ and CAGE^67^. The availability of high-quality genome annotations will further contribute to the effectiveness of these libraries.

Using PI-seq, we identified temporally resolved promoter activations during root colonization, including those associated with chemotaxis, plant immune suppression, biofilm formation, and ROS response, consistent with previous genetic studies^14,15,46,47^. In addition, we uncovered previously unrecognized transcriptional changes, such as the late-stage activation of the *xdh* and *mliC* genes, which further enhance the mutualistic interactions between *P. simiae* WCS417 and Arabidopsis through ROS neutralization and lysozyme inhibition, respectively (**Fig.4**). Because plant responses and root exudates strongly influence bacterial states, future studies could examine how specific metabolites and signaling molecules regulate bacterial gene expression. For example, coumarins exuded from Arabidopsis roots dynamically regulate genes involved in motility and biofilm formation in *P. simiae* WCS417^68^.

Notably, we observed significant dynamics in promoter activity during the first 24 hours, highlighting this period as critical for successful bacterial adaptation to the root environment. By contrast, relatively little change was observed at later time points (>24 hours) in both phytagel and soil experiments, likely reflecting reduced metabolic activity as cells enter a dormant phase after depleting locally available plant-derived nutrients. While our experiments relied on whole root sampling, finer-scale resolution could provide additional insight into how bacterial gene expression is influenced by spatial variation as root systems develop. For example, root tips secrete large amounts of metabolites, which can locally boost bacterial activity^69^. Although Arabidopsis roots are too small and fragile for such spatial dissection, plants with larger root systems, such as Brachypodium, may be a suitable model for future studies.

Among all promoters, the strongest upregulation occurred for those driving PQQ-related systems (PS417_RS10765 and PS417_RS10720), showing 206-fold and 24-fold increases, respectively, at 3 hours of colonization (**Fig.3K**). Notably, PS417_RS10720 controls a downstream response regulator transcription factor (PS417_RS10715), which shares 86% sequence identity with AgmR (also known as PedR1) of *Pseudomonas aeruginosa* PAO1—a key regulator of PQQ biosynthesis^70^. PQQ is a known cofactor involved in sugar and alcohol oxidation, with the metabolic byproducts that contribute to plant immune suppression by modulating local pH^14^. PQQ can also function as a growth-promoting factor when directly applied to plants^71,72^, and it stimulates production of natural products such as antibiotics in a broad range of *Actinobacteria*^73^. These findings underscore the vital role of PQQ production in the rhizosphere and its potential impact on microbial community structure and plant physiology.

Although PI-seq requires further refinement, this method provides a valuable tool for identifying regulatory elements associated with host-microbe interactions, particularly in non-model bacteria. Given the diverse roles that bacteria play in host physiology—such as biofertilization, biostimulation, and biocontrol—there is ample opportunity to uncover new molecular mechanisms. This knowledge will be instrumental in advancing the use of biologics, as well as in developing new biosensors and genetic circuits that can mediate interactions between microbes and their hosts^74–77^.

## Acknowledgements

We thank Zhiying Zhao, Robert Evans, William Kauffman, Jan-Fang Cheng, Benjamin Cole, Yuko Yoshinaga, Vlastimil Novak, William Kauffman, Emory Chan and Jeffery Dangle for helpful discussions and technical suggestions. We also thank Joint Genome Institute, QB3 Genomics Center at UC Berkeley, and Azenta Life Sciences for sequencing T.H. acknowledges a JSPS overseas research fellowship from the Japan Society for the Promotion of Science. Lawrence Berkeley National Laboratory is managed by University of California for DOE under contract number DE-AC02-05CH11231 (T.H. S.Y., D.M., L.B., Y.Y.). Argonne National Laboratory is managed by UChicago Argonne, LLC for DOE under contract number DE-AC02-06CH11357 (GB). This work was funded by the secure biosystem design (S.Y., G.B., Y.Y.) and the bioimaging programs (T.H., G.B., Y.Y.) of the U.S. Department of Energy, Office of Biological and Environmental Research under Contract No. DE-AC02-05CH11231 and DE-AC02-06CH11357. This work (proposal: 10.46936/10.25585/60001279), conducted by the U.S. Department of Energy Joint Genome Institute (https://ror.org/04xm1d337), a DOE Office of Science User Facility, is supported by the Office of Science of the U.S. Department of Energy operated under Contract No. DE-AC02-05CH11231 (T.H., D.M., L.B., Y.Y.). This project was also supported in part by the U. S. Department of Energy, Office of Science, through the Biomolecular Characterization and Imaging Sciences Program, Office of Biological and Environmental Research, under FWP 39156 (T.H., S.Y., Y.Y., G.B.). Raman Microscopy work at the Molecular Foundry was supported by the Office of Science, Office of Basic Energy Sciences, of the U.S. Department of Energy under Contract No. DE-AC02-05CH11231.

## Author Contribution

T.H. and Y.Y. designed the study. T.H. performed the experiments, with contribution by S.Y., who prepared sequencing libraries and characterized mutant phenotypes. T.H. and Y.Y. developed the computational pipeline. D.M. performed protein structural analysis. L.B. contributed to a plasmid construction and provided DAP-seq data. Y.Y and G.B. supervised the study and acquired funding. T.H. and Y.Y. wrote the manuscript. All authors reviewed and approved the final version.

## Data availability

All sequencing data generated and analyzed in this study are available at NCBI Short Read Archive: https://www.ncbi.nlm.nih.gov/sra/PRJNA1221951. Custom scripts used for PI-seq analysis is deposited at https://github.com/tomoyahonda/CRAGE-RB-PI-seq. Supplementary data that support this study are provided as downloadable Excel files (**Supplementary Data 1-5**).

