## Supplementary Information for "CRAGE-RB-PI-seq enables transcriptional profiling of rhizobacteria during plant-root colonization"

**1. Strain information** **2**
**1.1 Bacterial strains** **2**
**1.2 Design of barcoded promoter library** **2**
**1.3 Library synthesis, assembly, transformation, and conjugation** **2**
1.4 Construction of fluorescent reporter strains 3
**2. Bacterial and plant growth** **4**
**2.1 Bacterial cell culturing in liquid media** **4**
2.2 Plant growth conditions 4
**2.3 Bacterial root colonization assay** **5**
**2.4 Fluorescent quantification from seedling plates** **5**
**3. Sequence library preparation** **6**
**3.1 Library preparation to associate promoters and barcodes** **6**
**3.2 Nucleic acid extraction from root samples** **6**
**3.3 Library preparation for barcode amplicon** **6**
**3.4 RNA-seq library preparation** **8**
**4. Sequence data analysis** **8**
**4.1 Promoter and barcode association** **8**
**4.2 Barcode quantification** **9**
4.3 Differential expression analysis 9
**4.4 Functional analysis** **9**
**5. Mutant phenotype characterization** **10**
**5.1 Bacterial competition assay for root colonization** **10**
**5.2 Lysozyme assay** **10**
**5.3 Oxidative stress assay** **10**
**6. Supplementary figures** **11**
**7. Supplementary tables** **18**
**8. Supplementary data information** **21**
**9. References** **22**

### 1. Strain information

#### 1.1 Bacterial strains

We based our study on *Pseudomonas simiae* WCS417, originally obtained from Dr. Corne Pieterse (Utrecht University). This strain was domesticated by integrating a landing pad, resulting in strain SB599, as detailed by Wang *et al* (2020)[1]. The barcoded transposon library was generated by Cole *et al* (2017)[2]. The individual insertion mutants were later isolated at the Joint Genome Institute. *E. coli* WM3064 was obtained from Dr. William Metcalf (University of Illinois).

#### 1.2 Design of barcoded promoter library

We used the annotated genome of *Pseudomonas simiae* WCS417 (NCBI: NZ_CP007637) for a promoter library design. Individual genes were categorized into three groups based on the distances to upstream genes: group I (≤30 bp), group II (31–139 bp), and group III (≥140 bp). We extracted the 140 bp regions immediately upstream of the start codon for all genes using Geneious software (Dotmatics). To facilitate cloning, 30 bp priming sites were added to both ends of the 140 bp sequences. The three groups of 200 bp DNA libraries, referred to as promoter pools, were then synthesized by Twist Bioscience.

For the DNA barcodes, we designed 23 bp sequences, with the first three nucleotides set to CGT, followed by GGA, AGG, or GAG, depending on the group (I, II, or III). The remaining 17 bp were randomized. The first three nucleotides were designed to distinguish different strains for future studies, while the second set corresponded to the groups I-III, allowing for clear identification between them. We added 30 bp priming sites to both ends of the barcode sequences for cloning purposes, and the three groups of 83 bp DNA libraries were synthesized by Integrated DNA Technologies. These are referred to as barcode pools.

#### 1.3 Library synthesis, assembly, transformation, and conjugation

The following procedures were conducted in parallel for group I-III libraries. The oligo pools for the promoter and barcode pools were resuspended to 10 ng/µl and 10 µM, respectively. dsDNA pools were generated via PCR, using 1 µl of promoter libraries, 2.5 µl of barcode libraries, and 2.5 µl of 10 µM 5'-forward primers. PCR reactions were performed for 7 cycles in a total volume of 50 µl using Q5 polymerase (NEB: M0493L). The amplified products were subsequently gel-purified.

For the conjugal vector, we used a modified version of pW26[1] (pW26_mod), which contains two BsaI cloning sites and sfGFP, flanked by two mutually exclusive lox sites (*lox2272, lox5171*). The vector was digested with BsaI (NEB: R3733S) overnight and dephosphorylated using a quick dephosphorylation kit (NEB: M0525S). The digested vector was gel-purified and Gibson assembly was performed using ~23 ng of the digested vector and ~12 ng of dsDNA pools in a 10 µl reaction volume with a master mix (NEB: E2621S).

The assembled plasmids were transformed into *E. coli* EC100 pir+ competent cells (Lucigen: ECP09500) for amplification: After diluting the plasmids 5-fold in water, 1 µl was gently mixed with 20 µl of competent cells on ice. Following electroporation, a small aliquot of the recovery cultures was diluted and plated to determine cloning efficiency, while 500 µl of the cultures were diluted in 3 ml LB and spread onto bioassay dishes (VWR: 73520-774) containing LB agar and 50 µg/ml apramycin. The coverage of all group I-III libraries was determined to be >200×, calculated by dividing the number of colony-forming units by the size of the designed library. Plasmid DNA was then extracted from the colonies using a midiprep kit (Promega: A2492).

The extracted plasmids were subsequently transformed into *E. coli* WM3064 competent cells to be used as the donor strain for conjugation. After transformation, the cells were recovered in LB supplemented with 0.3 mM diaminopimelic acid (DAP) for 1.5 hours at 30 °C, and spread onto bioassay dishes containing LB agar, 50 µg/ml apramycin, and 0.3 mM DAP. The following day, cells were scraped from the plates, resuspended in 10 ml LB medium, and prepared for conjugation.

Library integration into the chromosome of *P. simiae* WCS417 was performed using CRAGE via conjugation, as described by Wang *et al* (2019)[3] and Wang *et al* (2020)[1]. We used *P. simiae* strain SB599, which harbors a *Cre* recombinase gene and kanamycin-resistant marker ($Km^R$) flanked by two mutually exclusive lox sites (*lox2272, lox5171*) in the landing pad, as a recipient strain. The landing pad is located at nucleotide location 268,402 between locus tag PS417_RS26490 and PS417_26495 without disrupting any genes[1]. For recombination, both donor and recipient strains were grown overnight in LB medium at 28 °C. The next morning, the cultures were washed twice with LB medium and mixed at a 4:1 (donor:recipient) ratio based on $OD_{600}$ to a total volume of 1 ml. Then the mixed cultures were washed once again and resuspended in 100 µl of LB + 0.3 mM DAP medium. The mixture was plated onto LB agar plates containing 0.3 mM DAP and incubated overnight at 28 °C.

The following day, all colonies were collected using a loop, resuspended in 1 ml LB medium, washed once, and spread onto bioassay dishes containing LB agar and 50 µg/ml apramycin. One day later, colonies were scraped from the plates, resuspended in 10 ml LB containing 10 % glycerol, and stored at -80 °C for later experiments.

#### 2. Bacterial and plant growth

##### 2.1 Bacterial cell culturing in liquid media

A 10 µl glycerol stock containing each group I-III library was inoculated into 3 ml fresh LB medium supplemented with 100 µg/ml apramycin and grown to saturation at 30 ℃ in a shaking incubator at 200 rpm. After pooling the group I-III cultures in equal ratios, 30 µl of the cultures were transferred into 3 ml of fresh M9-based growth media containing different nutrients and grown overnight. The following day, mid-log phase cultures were diluted to an $OD_{600}$ = 0.05-0.1 in pre-warmed growth media and cultured again at 30 °C (or 37 °C for heat stress conditions).

When the $OD_{600}$ reached ~1.0 (~$8x10^8$ cells/ml), 2 ml of the cultures were pelleted, and the supernatant was removed. The pellets were resuspended in 750 µl DNA/RNA Shield (Zymo Research: R1100), and the cells were lysed for 10 minutes using 0.1 mm and 0.5 mm beads at maximum speed. DNA and RNA were extracted in parallel using the ZymoBIOMICS DNA/RNA Miniprep kit (Zymo Research: R2002).

The growth media were based on M9 minimal medium, containing 1x M9 minimal salts (Gibco: A1374401), 2 mM magnesium sulfate, 0.1 mM calcium chloride, and 10 µM ferrous sulfate. The primary carbon sources used were 20 mM glucose, 40 mM glycerol, or 20 mM citrate. For high osmolarity conditions, sodium chloride was added to a final concentration of 300 mM.

##### 2.2 Plant growth conditions

*Arabidopsis thaliana* Columbia-0 (Col-0) seeds were surface-sterilized in 70 % ethanol for 5 minutes, followed by treatment with 50 % bleach plus 0.1 % Triton-X100 for another 5 minutes. The sterilized seeds were washed 5 times with sterile water and stratified in the dark for at 4℃ for 2-4 days. After stratification, approximately 100 seeds were plated on a nylon mesh filter (100 µm pore size [Genesee Scientific: 57-103], cut to approximately 8 $cm^2$) placed on top of plant growth media solidified with phytagel. The media consisted of 0.5 x Murashige and Skoog basal salts (PhytoTech Labs: M404), 2.5 mM MES (Sigma-Aldrich: M3671), and 0.6% phytagel (Sigma-Aldrich: P8169), with the pH adjusted to 5.7 using potassium hydroxide (Sigma-Aldrich: 319376), poured in a 10 cm square petri dish (Carolina Biological supply Company: 741470). The phytagel plates were sealed with micropore surgical tape (VWR: 56222-182) and grown upright in a Percival incubator (Geneva Scientific: CU36L5) for 10 days under long-day mode (16 hours light and 8 hours dark cycle) at 22 ℃ until exposure to bacterial cells.

##### 2.3 Bacterial root colonization assay

A 10 µl glycerol stock of the group II and III promoter libraries was inoculated into 3 ml of fresh LB medium supplemented with 100 µg/ml apramycin and grown for 5–6 hours at 28 °C in a shaking incubator (200 rpm) until the late exponential phase. Equal volumes of group II and III cultures were then pooled, and 3 µl of the mixed culture was transferred into 3 ml of M9 medium containing 20 mM glucose as the carbon source. After overnight incubation to mid-log phase, cells were harvested by centrifugation (3,000 g, 1 min), washed twice by pelleting and resuspension in 0.5× MS liquid medium, and finally resuspended to an OD600 of 0.5.

For phytagel plate experiments, 50 µl of the suspension was spread onto fresh phytagel plates using 5–10 sterile glass beads. Ten-day-old Arabidopsis seedlings grown on nylon mesh filters were transferred onto the plates using sterile tweezers. For clay experiments, 200 µl of the suspension was applied directly to the rhizosphere of Arabidopsis transplanted from germination plates into 130 g of clay supplied with 110 ml of 0.5× MS liquid medium in a vented vessel (PhytoTech Labs: C2110). Details of the clay substrate are described in Sasse *et al*[4].

Plates and vessels were incubated upright in a Percival incubator at 22 °C under a 16-hours light and 8-hour dark cycle. Root samples were collected at 10 min, 3 h, 24 h, and up to 7 days for phytagel experiments, and at 7 days and 21 days for clay experiments, following bacterial inoculation.

##### 2.4 Raman microscopy

Wild-type and group I–III library strains were pre-cultured as described in **Section 2.1**. After overnight growth, 1.4 ml of cell culture was collected, washed twice with sterile water, and resuspended in 20 µl of sterile water. An aliquot (1.5 µl) of the suspension was spotted onto quartz slides (Ted Pella: 26012) and air-dried at room temperature for 40 min. Raman spectra of dried cells were collected using a Horiba Jobin Yvon LabRAM ARAMIS confocal Raman microscope equipped with a 532 nm excitation laser (50 mW; Laser Quantum), a 532 nm long-pass filter (Semrock), a 1200 grooves/mm grating, a 200 µm pinhole, and a 100× 0.9 NA objective (Olympus). To minimize photodamage and ensure representative sampling, 285 spectra per sample were acquired by scanning the laser spot across a 19 × 15 pixel grid, with a pitch of 3 µm between pixels and an acquisition time of 1 s per pixel. Principal Component Analysis (PCA) was applied, when necessary, to distinguish spectra corresponding to cells from those lacking signal or corresponding to the quartz substrate. Cell spectra were then averaged and are presented without additional processing or background subtraction.

#### 3. Sequence library preparation

##### 3.1 Library preparation to associate promoters and barcodes

A 10 µl glycerol stock containing each group I-III library was inoculated into 3 ml of fresh LB media supplemented with 100 µg/ml apramycin and grown overnight at 30℃ in a shaking incubator at 200 rpm. After overnight growth, 300 µl of the saturated cultures were pelleted, and the supernatant was removed. The pellets were resuspended in 750 µl DNA/RNA Shield, and the cells were lysed for 10 minutes using 0.1 mm and 0.5 mm beads at maximum speed. DNA was extracted from each group I-III sample using a miniprep kit (Zymo Research: R2002).

For the first PCR, a region spanning the promoter to the barcode was amplified using specific primers (**Table S3**: map_fwd/rev). The same PCR settings as described below for the barcode amplifications (**section 3.3**) were used, except the number of cycles was set to 15. A second PCR was then performed to add indexes and Illumina P5 and P7 adaptors, following the same procedure as in **section 3.3**. The resulting amplicons were sequenced on Illumina NovaSeq platforms using 250 bp paired-end sequencing.

##### 3.2 Nucleic acid extraction from root samples

DNA and RNA were extracted from root samples using the ZymoBIOMICS DNA/RNA Miniprep Kit (Zymo Research: R2002) with a modified protocol to ensure efficient isolation and lysis of bacterial cells. To facilitate this, two different sizes of bashing beads were used. Seedlings were cut below the root/shoot junction, and the isolated roots were placed into 2 ml tubes. Roots from one plate (30-50 seedlings, ~20 mg) were pooled into a single sample.

The pooled roots were vortexed for 5 seconds in 800 µl of M9 buffer to remove loosely adhered cells (and clay when used) from the root surface. After the buffer was removed, 800 µl of DNA/RNA Shield and 2 mm beads (Zymo Research: S6003-50) were added to the tubes. The tubes were then placed onto an adapter (Zymo Research: S5001-7) attached to a Vortex Genie2, and the samples were ground for 10 minutes at maximum speed. Once the root tissues were disrupted, the lysed samples were transferred to tubes containing 0.1 mm and 0.5 mm beads from the R2002 kit and ground for another 40 minutes at maximum speed to lyse the bacterial cells. Following this, we used the parallel DNA and RNA extraction procedure, including DNAse treatment, as described in the manufacturer's protocol (Zymo Research: R2002). DNA and RNA were eluted in 70 µl of ddH$_2$O and stored at -20 ℃.

##### 3.3 Library preparation for barcode amplicon

To create sequencing libraries, barcoded regions of genomic DNA or cDNA were amplified in a two-step PCR process. For genomic DNA samples, DNA was amplified using primers flanking the barcode region (**Table S3**: barcode_fwd/rev). Six tubes of 50 µl reactions were prepared for each sample, with the following components:

- 10 µl of Q5 reaction buffer (New England Biolabs: B9027S)
- 1 µl of dNTP (New England Biolabs: N447L)
- 2.5 µl of 10 µM forward primer (barcode_fwd)
- 2.5 µl of 10 µM reverse primer (barcode_rev)
- 10 µl of purified DNA
- 0.5 µl of Q5 polymerase (New England Biolabs: M0493L)
- 10 µl of betaine solution (Sigma-Aldrich: B-0300)
- 13.5 µl of ddH$_2$O

The PCR conditions were as follows:
1) initial heating at 98 ℃ for 30 seconds
2) 14 cycles of 10 seconds at 98 ℃ and 60 seconds at 72 ℃
3) A final extension at 72 ℃ for 60 seconds.

The six reactions were pooled into a single sample and purified using the DNA Clean & Concentrate Kit (Zymo Research: D4013), eluted into 20 µl of ddH$_2$O, and quantified using a Qubit fluorometer (Thermo Scientific).

For the second PCR, we added indexes and Illumina P5 and P7 adaptors to the samples. A 50 µl reaction was prepared as follows:

- 10 µl of Q5 reaction buffer (New England Biolabs: B9027S)
- 1 µl of dNTP (New England Biolabs: N447L)
- 2.5 µl of 10 µM indexed P5 adapter primer
- 2.5 µl of 10 µM indexed P7 adapter primer

- ~30 ng of diluted DNA samples
- 0.5 µl of Q5 polymerase (New England Biolabs: M0493L)
- $ddH_2O$ to a total volume of 50 µl

The PCR conditions were as follows:
1) Initial heating at 98℃ for 30 seconds
2) 5 cycles of 10 seconds at 98 ℃, 15 seconds at 60 ℃, and 30 seconds at 72℃
3) A final extension at 72 ℃ for 60 seconds.

The samples were then gel-purified and eluted into 30 µl of $ddH_2O$ using a cleanup kit (Macherey-Nagel: 740609.050). The quality of the amplicon samples was validated using Bioanalyzer DNA Analysis (Agilent: 5067-1504).

For RNA samples, 70 µl of purified RNA was treated with DNase (Thermo Scientific: AM1907) to remove any trace genomic DNA. RNA quality was checked using Bioanalyzer RNA Analysis (Agilent: 5067-1511), and samples were concentrated with the RNA Clean & Concentrate Kit (Zymo Research: D1013). Selective reverse transcription was carried out using SuperScript II Reverse Transcriptase (invitrogen: 18064022) with a 1st strand synthesis primer (**Table S3**: cDNA_syn), following the manufacturer's protocol. Synthesized cDNA was amplified by PCR in eight tubes of 50 µl reactions, each containing the same components as for genomic DNA amplification, except using 2 µl of cDNA and 21.5 µl of $ddH_2O$. The PCR conditions were as follows:
1) Initial heating at 98 ℃ for 30 seconds
2) 22 cycles of 10 seconds at 98 ℃ and 60 seconds at 72 ℃
3) A final extension at 72 ℃ for 60 seconds.

The eight reactions were pooled, purified using the DNA Clean & Concentrate Kit (Zymo Research: D4013) and eluted in 20 µl of $ddH_2O$. The quality of the PCR products was validated using Bioanalyzer High-sensitivity DNA Analysis (Agilent: 5067-4626). Sequencing libraries were prepared by adding indexes and Illumina P5 and P7 adaptors as described earlier.

For samples from liquid cultures, the cycle numbers of the first PCR were reduced to 12 for genomic DNA and 20 for cDNA. Otherwise, the same procedures were followed. Amplicon samples were sequenced on Illumina NovaSeq platforms using 150 bp paired-end sequencing, targeting 10-20 million reads per sample.

##### 3.4 RNA-seq library preparation

For samples from liquid culture, ribosomal RNA (rRNA) was removed from 1 µg of total RNA using an rRNA depletion kit (New England Biolabs: E7860S), and the treated RNA was eluted in 6.5 µl of $ddH_2O$. Sequencing libraries were prepared using the TruSeq Stranded mRNA Library Prep kit (Illumina: 20020594). This procedure involved adding 3 µl of rRNA-depleted RNA into 15 µl of fragment, prime, and finish (FPF) buffer, followed by the manufacturer's protocol. Library quality was validated using Bioanalyzer DNA Analysis (Agilent: 5067-1504). Samples were sequenced on Illumina NovaSeq platforms using 150 bp paired-end sequencing, targeting 5 million reads per sample.

For RNA samples from Arabidopsis seedlings, rRNA depletion was not performed, as our goal was to determine the fraction of reads mapped to Arabidopsis and *P. simiae* (**Table S2**). We used 60 ng of total RNA, starting the procedure by adding the RNA to 18 µl of FPF buffer, then following the same steps as described above. Samples were sequenced on Illumina NovaSeq platforms using 150 bp paired-end sequencing, targeting 400 million reads per sample.

#### 4. Sequence data analysis

##### 4.1 Promoter and barcode association

A custom shell script using BBtools (https://sourceforge.net/projects/bbmap/), along with an R script incorporating the Biostrings (https://bioconductor.org/packages/Biostrings) and dplyr (https://dplyr.tidyverse.org/) packages, was used to map barcode sequences to their corresponding promoter sequences. Briefly, 140 bp promoter regions and 20 bp barcodes (excluding initial CGT specific to *P. simiae* WCS417) were extracted from read 1 sequences. Promoter-barcode paris were retained if they were detected more than 100 times in the sequencing reads. Barcodes mapped to two or more distinct promoter sequences were excluded from further analysis. Promoter sequences that exactly matched their native counterparts were then selected. These steps were repeated for each group (I-III). Overall, 91% (5,040 promoters) of the total 5,541 promoters were successfully assigned unique barcodes, with each promoter associated with an average of 50 unique barcodes.

##### 4.2 Barcode quantification

Barcode sequences were extracted and counted from the sequencing data using a custom shell script with BBtools and an R script utilizing Biostrings and dplyr packages. Barcode counts were normalized to sequencing depth for each sample and then summed for each promoter. Library coverage was assessed by normalizing DNA barcode counts to counts per million (CPM). To calculate individual promoter activity (in arbitrary units, A.U.), the barcode counts from RNA-derived cDNA were divided by the barcode counts from genomic DNA.

##### 4.3 Differential expression analysis

For the plant assay, statistical analysis was carried out using edgeR and limma packages, with custom modifications based on Law *et al* (2018)[5]. Briefly, raw barcode counts were normalized using the trimmed mean of M values (TMM) and transformed to $\log_2$ counts per million ($\log_2$-CPM) with associated precision weights using voom. Linear models were then fitted for each promoter, and empirical Bayes moderation was applied with the trend and robust options enabled. Differential activity was quantified as $\log_2$ fold-changes with associated p-values, and p-values were adjusted for multiple testing using the Benjamini–Hochberg method. For the liquid culture experiments, differentially expressed promoters were selected based on $|Log_2FC| > 2.5$, with inactive promoters ($log_2$ promoter activity < 2) filtered out. In addition, promoters with zero RNA barcode counts in glucose conditions were excluded from the analysis.

##### 4.4 Functional analysis

Operon structures were predicted using Rockhopper[6], using the *P. simiae* WCS417 genome sequence and RNA sequencing reads from five different liquid cultures as input. Before analysis, RNA reads mapped to rRNA were filtered out using BBsplit function from BBtools. KEGG pathway analysis was performed using the kegga function from the limma package[7] in R. For promoters identified as driving operons (based on the Rockhopper results), clusters of genes driven by the promoter were used as input for pathway analysis. KEGG IDs of the genes were obtained from IMG[8].

##### 4.5 RNA-seq data analysis

RNA-seq data were processed using a custom shell script with BBtools and R scripts using dplyr, edgeR and limma packages. In this pipeline, reads mapped to 5s, 16s, and 23s rRNA were filtered out using the BBsplit function. For root samples, Arabidopsis genes were also filtered out. The remaining reads were mapped to the *P. simiae* WCS417 genome using HISAT2[9]. Mapping results were used to calculate gene-level read counts with FeatureCounts[10]. These count data were normalized to CPM for each sample, and differentially expressed genes were identified based on a $|Log_2FC| > 2$.

#### 5. Mutant phenotype characterization

##### 5.1 Bacterial competition assay for root colonization

Wild-type and individual mutant strains (kanamycin-resistance) were cultured as described in **section 2.3**, using 1x M9 minimal medium with 20 mM glucose (no antibiotics) for pre-cultures, and 1x M9 minimal salts (Gibco: A1374401) for washing. Cell cultures were normalized to an $OD_{600}$ of 0.5, and wild-type and mutant strains were mixed at a 1:1 ratio. A 50 µl aliquot of this mixture was spread onto phytagel plates. 10-day-old Arabidopsis seedlings, grown on nylon mesh filters, were then transferred onto the inoculated plates using sterile tweezers. The plates were incubated upright in a Percival incubator at 22 ℃ with a 16-hour light and 8-hour dark cycle.

After 5 days of incubation, root samples were collected and washed once with 1x M9 buffer. The buffer was removed, and 800 µl of fresh buffer along with 2 mm beads (Zymo Research: S6003-50) were added to the tubes. The tubes were placed on a vortex adapter, and the roots were ground at maximum speed for 10 minutes. The lysed samples were diluted 10,000-fold, and 100 µl were spread onto LB plates containing 200 µg/ml kanamycin. After incubation at room temperature for two days, colony numbers were determined. Mutant cell counts were calculated based on colonies growing on LB plates with kanamycin, while wild-type counts were determined by subtracting the mutant colony numbers from the colony numbers on LB plates. Typically, around 20 mg of root tissues was collected per experiment.

#### 5.2 Lysozyme assay

To investigate the function of the MliC family protein (PS417_RS02110), we tested the sensitivity of both the wild-type and the mutant strain to lysozyme. Cells were cultured in LB medium and then transferred to M9 minimal medium with 20 mM glucose at 28°C. After overnight incubation, cells were washed three times with phosphate-buffered saline (PBS) and resuspended to an $OD_{600}$ of 0.05. Lysozyme (Roche: 10837059001) was then added to a final concentration of 25 mg/ml. Control samples were incubated in PBS without lysozyme. After 24 hours of incubation, cells were serially diluted and plated onto LB agar plates. Colony counts were determined after 48 hours of incubation at 28°C.

#### 5.3 Oxidative stress assay

The sensitivity of both wild-type and mutant strains of xanthine dehydrogenase (PS417_RS04040) to hydrogen peroxide ($H_2O_2$) was determined by monitoring growth curves. Cells cultured overnight in LB medium were transferred into M9-glucose medium at a 1:1000 dilution and incubated at 28 °C overnight. The following day, cells were transferred to a 96-well plate containing 200 µL of M9-glucose medium, adjusted to a starting $OD_{600}$ of 0.1. The cells were grown in a microplate spectrophotometer (Infinite 200 PRO, Tecan, Switzerland) with absorbance at 600 nm measured every 30 minutes. $H_2O_2$ (final concentration of 5 mM) was added after 7 hours of incubation, which was set as time 0 in **Fig.4D**.

#### 6. Supplementary figures

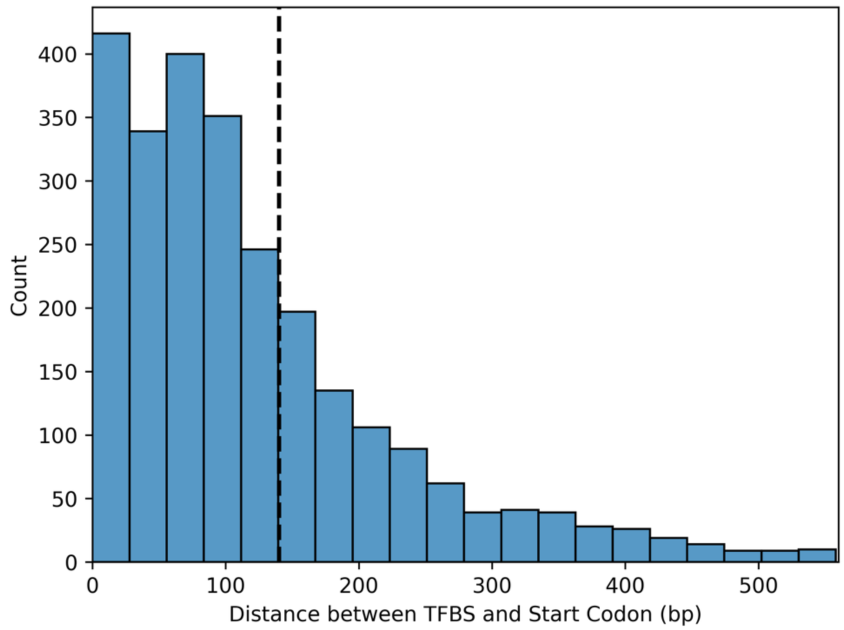


**Supplementary figure 1: Histogram of distances between TF binding sites and start codons.** Histogram showing the distance between transcription factor binding sites (TFBSs) identified in Baumgart *et al*[11] and the start codon of the immediately flanking gene(s) in *P. simiae* WCS417. Only significant binding sites with at least 15-fold enrichment were considered. 66% of TFBSs fall within the first 140 bp directly upstream of start codons. Dotted line is shown at 140 bp, corresponding to the promoter size used in the current study.

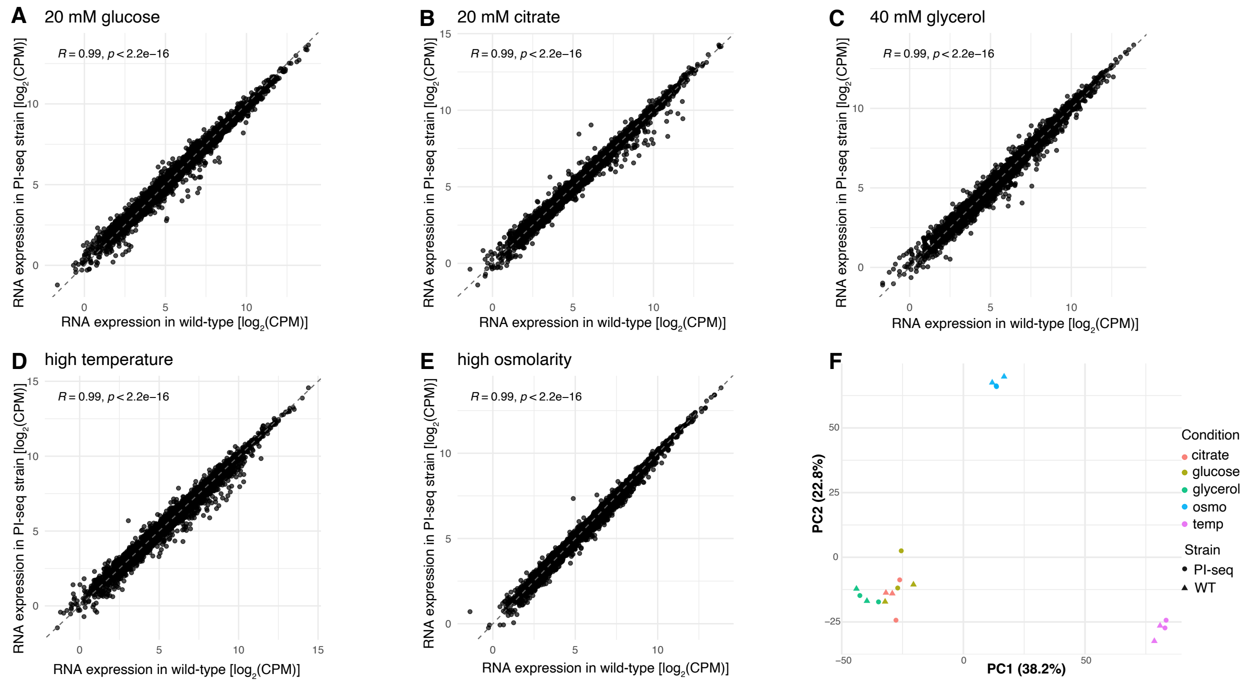


**Supplementary figure 2: Comparison of wild-type and engineered strains by transcriptome analysis.** Wild-type and promoter library strains (Groups I–III) were cultured under five conditions (A–E). Transcriptomes were measured by RNA-seq, revealing high correlations between the two strains (n = 2 biological replicates). The PCA plot in (F) further demonstrates clear clustering of the two strains.

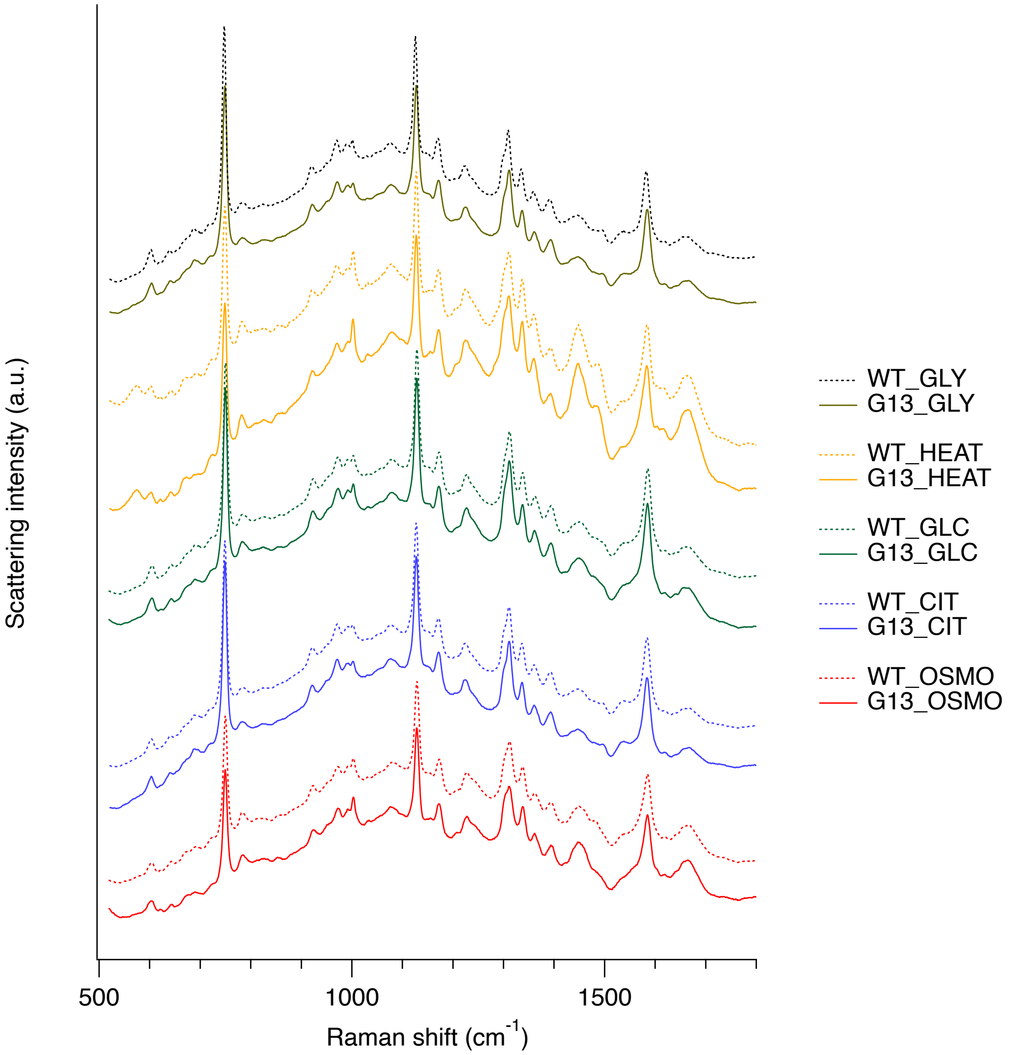


**Supplementary figure 3: Comparison of wild-type and engineered strains by Raman microscopy.** Wild-type and promoter library strains (Groups I–III) were cultured under five conditions as listed. Raman spectra were acquired and averaged across 285 measurements per sample, revealing highly similar peak profiles between the two strains.

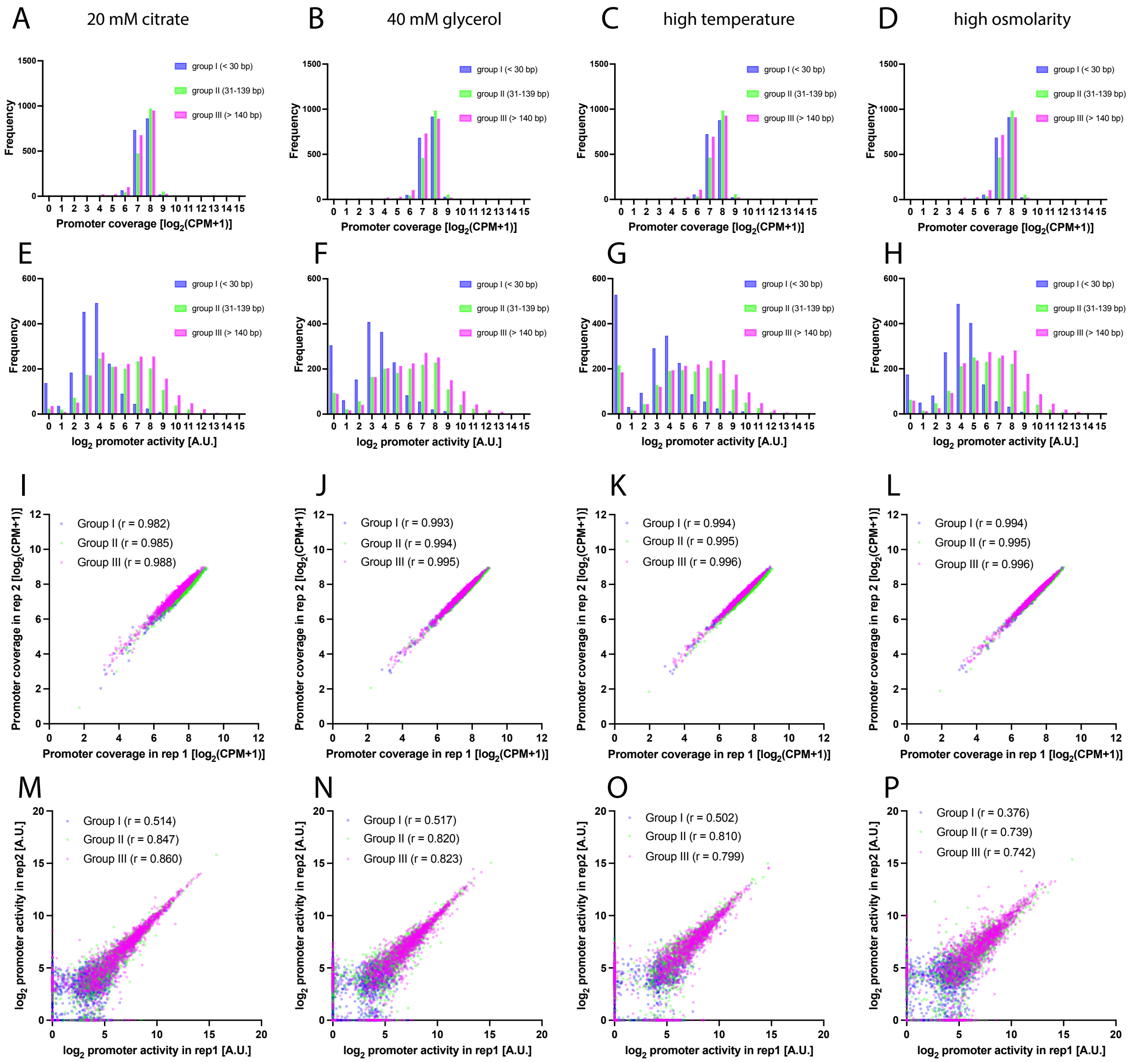


**Supplementary figure 4: Characterization of promoter library coverage and promoter activity under various liquid media.** In addition to the data presented in Figure 2 (20 mM glucose), we further analyzed PI-seq data for the cells grown under 4 additional culture conditions (*n* = 2 biological replicates). The results, displayed from left to right, show the data from 20 mM citrate, 40 mM glycerol, 20 mM glucose at 37 °C (high temperature), and 20 mM glucose with 300 mM NaCl (high osmolarity). These data demonstrate the high coverage of the promoter library and reproducibility of promoter activity measurements across conditions. (**A-D**) Histograms displaying promoter library coverages, based on DNA barcode counts, show consistent distribution across all conditions (**E-H**) Histograms of promoter activities, determined by normalizing RNA barcode counts with corresponding DNA barcode counts, indicate a large variation in promoter activity levels (**I-L**) Scatterplots showing the reproducibility of promoter coverages between two biological replicates (**M-P**) Scatterplots showing the reproducibility of promoter activities between two biological replicates, further supporting the reliability of the PI-seq data.

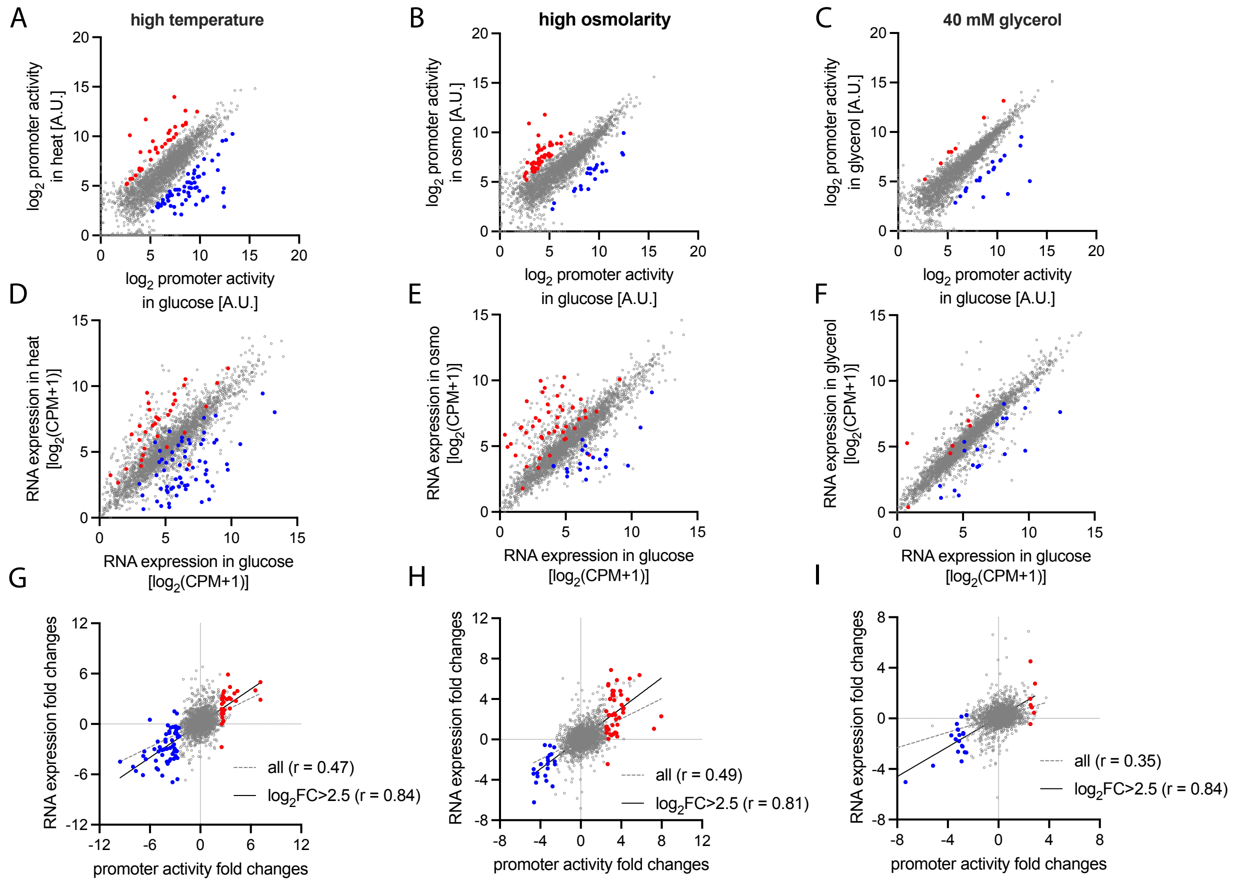


**Supplementary figure 5: Comparison of PI-seq to RNA-seq in other culture conditions.**
In addition to the data shown in Fig.2, we compared PI-seq data with RNA-seq data for cells grown in three additional culture conditions. Each column (from left to right) represents comparisons between data obtained in the 20 mM glucose condition and those from heat stress (left), osmotic stress (middle), and 40 mM glycerol (right). (**A-C**) Scatterplots of PI-seq data, comparing promoter activity between 20 mM glucose and each individual condition. Promoters upregulated in glucose and the respective conditions are highlighted in blue and red, respectively. Differentially expressed promoters were selected based on a $|Log_2FC| > 2.5$, after filtering out low-activity promoters ($log_2$ promoter activity < 2). (**D-F**) Scatterplots of RNA-seq data, comparing gene expression between 20 mM glucose and each condition. Genes driven by the promoters identified in PI-seq (A-C) are highlighted in corresponding colors. (**G-I**) Scatterplots comparing fold changes in promoter activity (x-axis, PI-seq) with RNA expression (y-axis, RNA-seq). Fold changes were calculated by dividing expression values in each condition by those in 20 mM glucose. Pearson's r correlation values are provided for all genes (dashed gray) and for differentially expressed genes ($|log_2FC| > 2.5$, solid black). The data of comparative analysis of in vitro culture conditions between PI-seq and RNA-seq is provided in **Supplementary Data 3**.

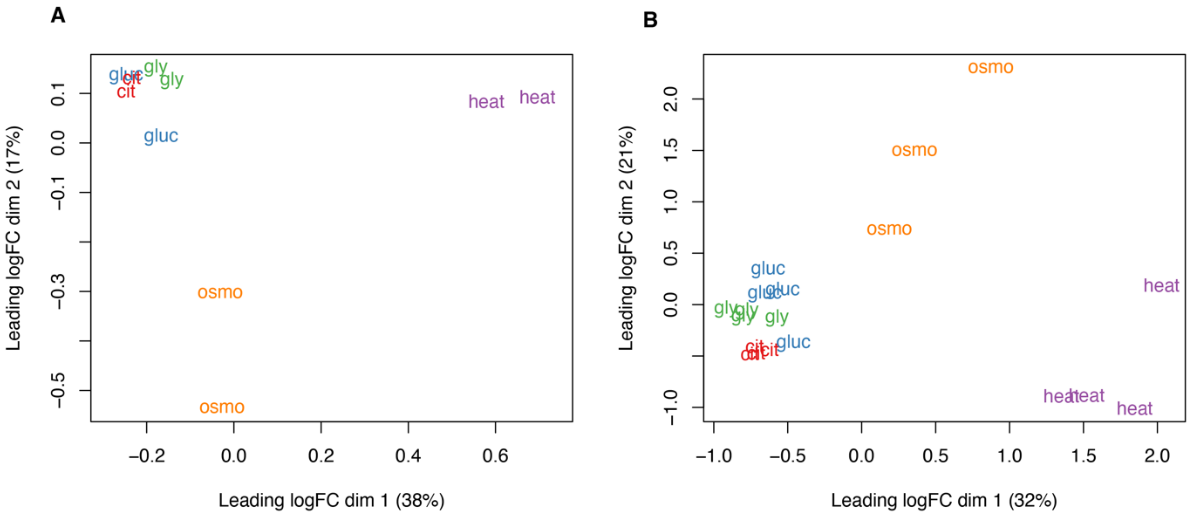


**Supplementary figure 6: Unsupervised clustering of PI-seq and RNA-seq samples.** Multi-dimensional scaling (MDS) plots of PI-seq (A) and RNA-seq (B) data from various liquid culture conditions. Both PI-seq and RNA-seq data show similar clustering patterns, where samples from different nutrient conditions: 20 mM glucose (gluc), 20 mM citrate (cit), 40 mM glycerol (gly) cluster together, while heat and osmotic stress conditions form distinct clusters.

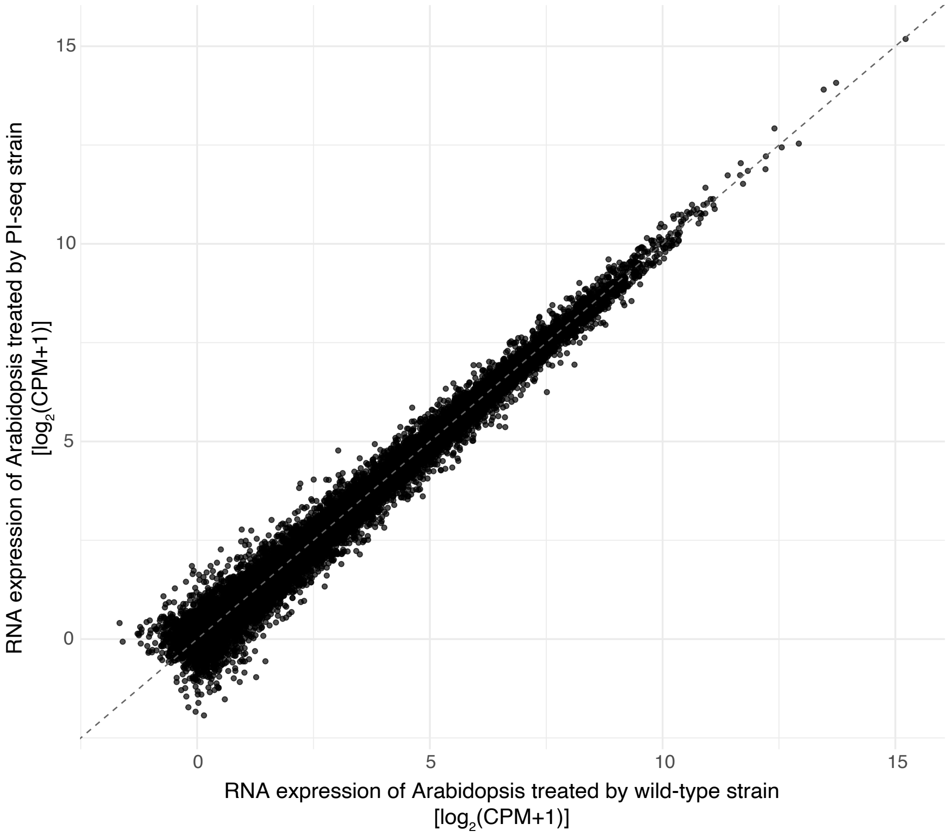


**Supplementary figure 7: Comparison of Arabidopsis root transcriptomes after inoculation with wild-type and PI-seq *P. simiae* strains.** RNA-seq analysis of Arabidopsis roots 4 hours after inoculation with either wild-type or PI-seq *P. simiae* strain. Three biological replicates were analyzed per condition. Differential expression analysis identified no significantly differentially expressed genes between treatments, indicating that the engineered strain does not alter the Arabidopsis transcriptional response compared to the wild type.

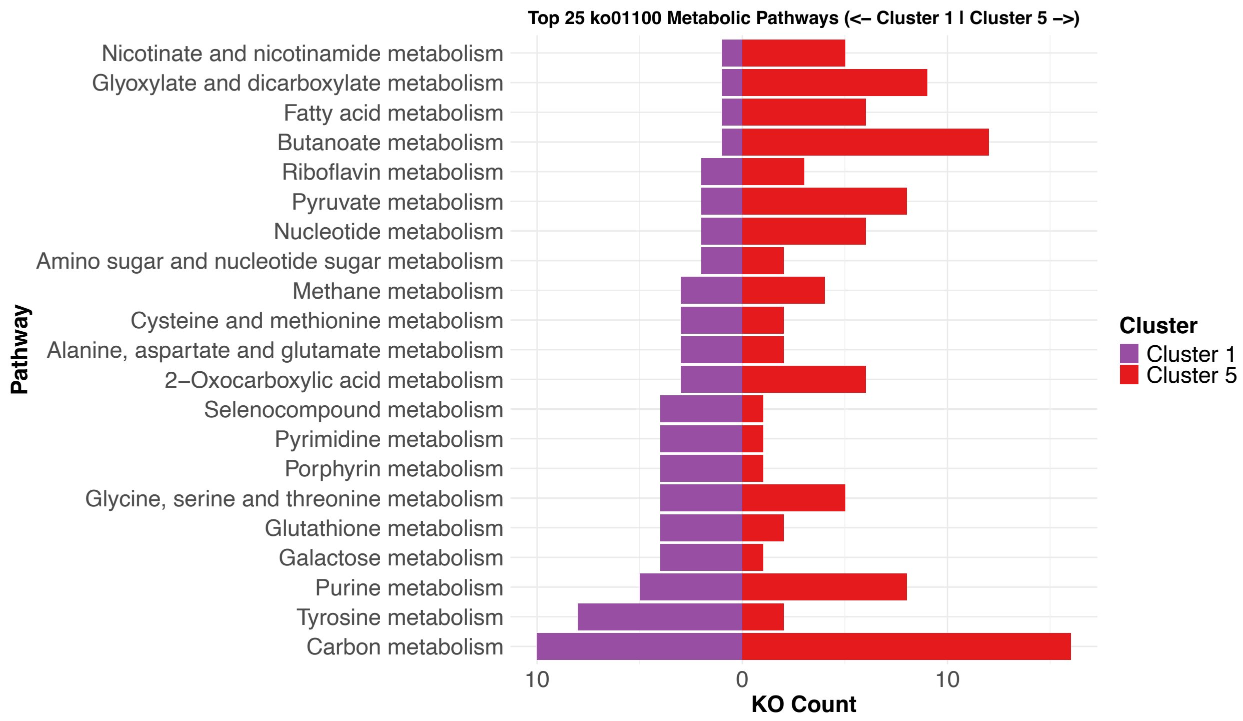


**Supplementary figure 8: Comparison of activated metabolic pathways in cluster 1 and 5.** Genes assigned to the KEGG category “metabolic pathways” in clusters 1 and 5 were further examined to identify the specific pathways involved, revealing distinct profiles between the two clusters.

A

| Query against PS417_RS02110 | Protein sequence identity (%) | Structural alignment (RMSD) |
|---|---|---|
| *E.coli* K12 *mliC* | 32/111 (29%) | 0.946 |
| PA0867 \| *mliC* | 18/77 (23%) | 1.943 |

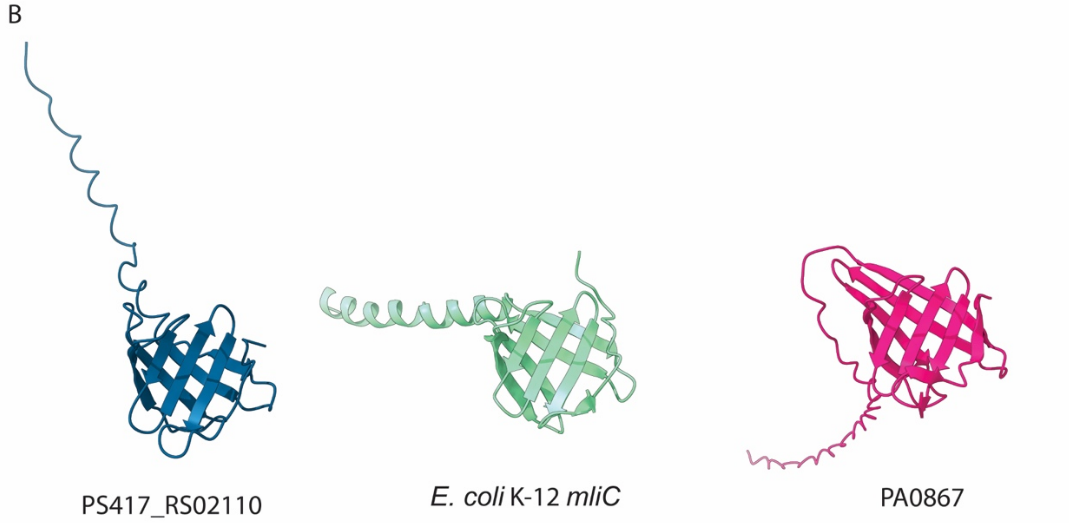


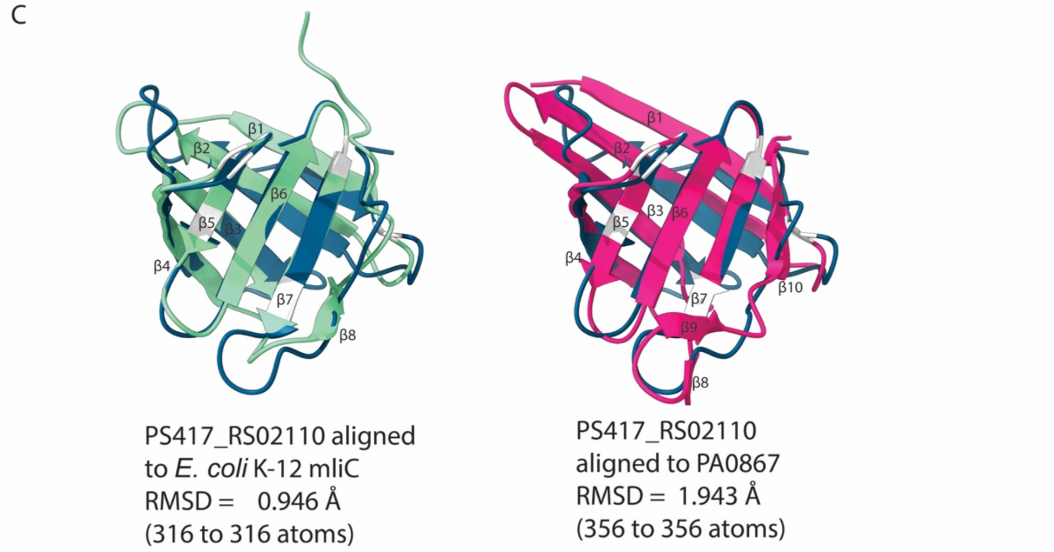


**Supplementary figure 9: Sequence and structure analysis of MliC.**

(**A**) Summary of protein sequence identity and structural alignment of the MliC family proteins using pairwise alignment and AlphaFold 3. MliC from *P. simiae* WCS417 (PS417_RS02110) was compared against the homologs in *E. coli* K12 and *Pseudomonas aeruginosa* (PA0867). (**B-C**) Predicted structures of MliC for each species, generated by AlphaFold 3, along with their structural alignment to assess similarities.

#### 7. Supplementary tables

| Strain | Parent* | Description | Antibiotics |
|---|---|---|---|
| Promoter library group I | SB599 | Promoters selected from the genes whose intergenic distances towards upstream genes are ≤30 bp | Apramycin 50ug/ml |
| Promoter library group II | SB599 | Promoters selected from the genes whose intergenic distances towards upstream genes are 31-139 bp | Apramycin 50ug/ml |
| Promoter library group III | SB599 | Promoters selected from the genes whose intergenic distances towards upstream genes are ≥140 bp | Apramycin 50ug/ml |

**Table S1: Strains constructed in this study.**

*The parental strains SB599 was constructed by Wang et al (2020)[1]

| | Total reads | *Pseudomonas simiae* WCS417r | | *Arabidopsis thaliana* |
|---|---|---|---|---|
| | | 5s, 16s, 23s rRNA | CDS | |
| **Replicate 1** | 452,547,206 | 26,375,444 (5.83 %) | 211,466 (0.05 %) | 424,401,754 (93.78 %) |
| **Replicate 2** | 397,440,972 | 46,299,996 (11.65%) | 280,460 (0.07 %) | 349,387,168 (87.91 %) |
| **Replicate 3** | 431,512,824 | 32,709,114 (7.58 %) | 214,280 (0.05 %) | 397,143,752 (92.04 %) |

**Table S2: Read counts mapped to *P. simiae* and Arabidopsis from RNA-seq data.** RNA-seq libraries were prepared from RNA extracted from Arabidopsis seedling roots colonized by *P. simiae* on sampling day 1 ($n$ = 3 biological replicates). The table summarizes the number of reads mapped to the *P. simiae* and Arabidopsis genomes.

| Name | Sequence |
| --- | --- |
| barcode_fwd | CTCTTTCCCTACACGACGCTCTTCCGATCTCTGTGCCAAGCGAACTGAGTGTATTTGAGC |
| barcode_rev | GAGTTCAGACGTGTGCTCTTCCGATCGGCAGTTTACCAGTAGTACAGATGAACT |
| map_fwd | CTCTTTCCCTACACGACGCTCTTCCGATCTTTACTGGATCTATCAACAGGAGTCC |
| map_rev | GAGTTCAGACGTGTGCTCTTCCGATCGGCAGTTTACCAGTAGTACAGATGAACT |
| cDNA_syn | GGCAGTTTACCAGTAGTACAGATGAACT |

**Table S3: Primers used in this study.**

#### 8. Supplementary data information

**Supplementary data 1:**
A list of promoters with their assigned barcodes obtained from DNA-seq

**Supplementary data 2:**
A list of multi gene operons and single genes obtained from in silico operon structure prediction

**Supplementary data 3:**
Comparative analysis of in vitro culture conditions between PI-seq and RNA-seq

**Supplementary data 4:**
A list of regulated promoters and their encoding genes in root colonization on phytagel

**Supplementary data 5:**
A list of regulated promoters and their encoding genes in root colonization in clay
